# Enteric ChAT-expressing neurons as new target for *Lactobacillus plantarum* ameliorates inflammatory bowel diseases

**DOI:** 10.1101/2025.09.09.673902

**Authors:** Bing Xia, Yuwei Sun, Jialin Wang, Yike Liu, Lu Wang, Shiqi Jiang, Gaiyan Bai, Ruijing Zhang, Yu Zhang, Rongwei Kou, Danna Wang, Beita Zhao, Junjie Yi, Yu Fu, Xiaoning Liu, Xuebo Liu

## Abstract

The gut microbiota plays a crucial role in inflammatory bowel diseases (IBD), yet how specific microbial components influence disease progression remains incompletely understood. Our study reveals that decreased abundance of Lactobacillus species correlates with ulcerative colitis severity in patients. Among tested strains, *L. plantarum* A736, isolated from traditional fermented foods, demonstrated remarkable efficacy in ameliorating dextran sulfate sodium (DSS)-induced colitis in a mouse model. Strikingly, multi-omics analysis revealed that *L. plantarum* A736 uniquely enhances choline acetyltransferase-expressing (ChAT^+^) neurons in the enteric nervous system and elevates acetylcholine (ACh) levels. Further analysis identified that *L. plantarum* A736 metabolizes tryptophan to produce indole-3-lactic acid (ILA), and supplementation with ILA significantly mimics this strain’s anti-inflammatory effects by activating ChAT^+^ neurons. Crucially, selective ablation of colonic ChAT^+^ neurons completely abolished the therapeutic benefits of both *L. plantarum* A736 and ILA, establishing their essential role in mediating these effects. These findings demonstrate for the first time that microbial metabolites can directly modulate enteric neuronal circuits to regulate gut immunity. The identified ILA-ChAT^+^ neuron axis represents a novel neuro-immune pathway that could be targeted for IBD treatment.

**Graphical abstract:** 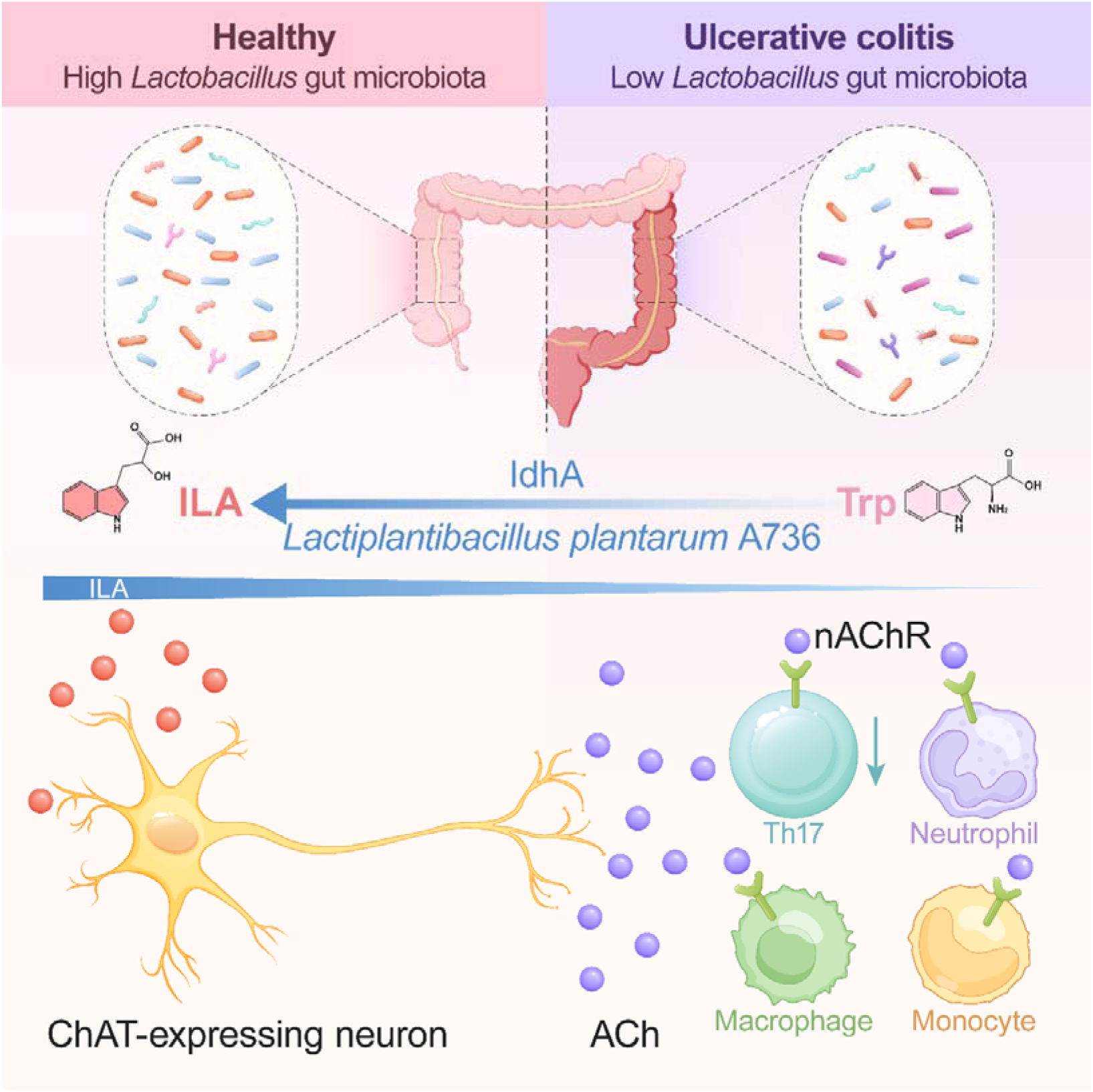

## Introduction

Inflammatory bowel disease (IBD), which primarily includes Crohn’s disease and ulcerative colitis (UC), is the most common chronic inflammatory disorders characterized by immune-mediated intestinal inflammation that is caused by a confluence of host genetics, environmental exposures and microbial factors.^1–3^ It has become a global disease and affects 6·8 million people worldwide,^4,5^ while the complex pathogenesis of disease remains unclear. Currently, available therapies targeting various inflammatory pathways cannot eventually cure IBD with a high a relapse and unexpected adverse events.^6^ Given the lack of treatment response, there is an urgent clinical need to make mechanistic dissection of involved pathways in diseases that reveal novel targets and further develop new effective treatments targeting specific pathways to break the therapeutic ceiling.

The enteric nervous system (ENS), known as ‘the first brain’ on presumed evolutionary grounds, is the intrinsic neuroglial circuits of the gastrointestinal tract containing neurons and enteric glial cells (EGC), which coordinates diverse aspects of intestinal physiology.^7–9^ Their disturbances of structure and function have been demonstrated with potential links to IBD.^8,10,11^ Indeed, risk genes for IBD, including BTBD8, GRP, JAZF1, NDFIP1, PLA2R1, and CNTNAP2, are enriched in human enteric neurons, suggesting neuronal contributions to disease.^12^ As observed in UC patients, the density of neurons is decreased in colon of mice with DSS-induced colitis.^8,10,13,14^ Another evidence indicates that pharmacologic ablation or chemogenetic inhibition of catecholaminergic axons has protective properties against colitis.^15^ Furthermore, cholinergic neurons are a major subset of enteric neurons expressing choline acetyltransferase (ChAT).^12,16^ Several seminal publications have extensively discovered the loss or dysfunctions of enteric cholinergic neurons in both patients and animal models with IBD.^14,17–21^ Upon activation, the nerve terminals of nociceptors release neurotransmitter acetylcholine (ACh), which are involved in manipulation of immune cell function through ACh receptors on immune cells, including mAChRs and nAChRs respectively. For example, the alleviation of colitis achieved by ACh supplementation attributes to activating the nAChR/ERK/IL-10 pathway in monocytic myeloid-derived suppressor cells (M-MDSCs).^14^ Furthermore, ACh exerts a strong effect on lipopolysaccharide (LPS)-activated macrophages to reduce TNF expression.^17^ The same phenomenon could be observed in mice treated with nicotine supplementation, due to the ability to restore nicotinic acetylcholine receptor signaling levels, which relieves colitis after DSS exposure.^22^ Because ENS is difficult to assess and analyze in humans and, often, enteric neurons are rare and diverse, further investigation is needed to improve understanding of ENS molecular mechanisms, which can constitute possible novel targets for the treatment of IBD.

It is generally accepted that inappropriate immune activation has been identified as a a key feature of IBD, although the cause of IBD is complicated and poorly understood.^23^ In colon lamina propria (LP), immune cells localize in proximity to nerve fibers of enteric neurons. Based on neurotransmitter, neuropeptide and cytokines released by enteric neurons and the expression of their receptors in immune cell subsets, enteric neurons can directly affect immune cell functions.^24^ The crucial function of neuroimmune interactions have been described as important pillars of intestinal physiology.^25^ Recently, multiple lines of studies have uncovered that ENS has an important role in modulating immune cells.^26^ For example, interleukin-6 released from enteric neurons tunes Treg cell generation Treg cell generation to affect intestinal tolerance.^27^ Another recent study has focused on enteric vasoactive intestinal peptide (VIP)-producing neurons, which modulates intestinal immune homeostasis through VIP receptor (VIPR2) on CCR6+ ILC3 cells^28^ inhibit IL-22 production. Further, two recent studies demonstrate that neuron-derived neuropeptide neuromedin U (NMU) controls group 2 innate lymphoid cells (ILC2s) which results in regulating tissue protection and inflammation.^29,30^ Beyond these one-to-one interactions, neuroimmune circuits remains challenging due to the diversity of enteric neuronal subtypes and immune cell populations. It remains an intriguing possibility that neuroimmune interactions might be the preferred route in IBD targets and strategies.

Many anatomical and cellular blueprints of the ENS are in place by birth, but ENS activity and function is primarily regulated by the luminal commensal microbiota.^16,24,28,31–34^ Alternatively, probiotics emerge as a means to modulate the gut microbiota.^35^ Much of our insights on probiotic mechanisms profiles mainly include the production of specific metabolites (such as bile acids, short-chain fatty acids and tryptophan metabolites) and direct interactions with host cells or the resident microbiota,^35^ although understanding of interactions with different intestinal cell types have not yet been fully defined. Also, the transcription factor aryl hydrocarbon receptor (Ahr), the activity of which is regulated by microbial-derived notably tryptophan metabolites, is expressed predominantly in myenteric neurons.^31^ This suggests probiotics as a potential modulatory tool to influence ENS function. Accumulated evidence on both animal and clinical studies have indicated that probiotics are microbiota-management tools for modulating IBD. As such, a report from the American Gastroenterological Association Institute also suggested that p_r_obiotics may be considered for treatment of functional symptoms in IBD.^36^ Probiotics can also emerge as adjuvant approach for IBD by modulating gut microbiota.^37,38^ Thus, the mechanistic basis for probiotics involved in the prevention of IBD needs to be better explored/ remains to be gathered compared to present drugs. Recently, one group notes an increase in the number of enteric neurons of IBS mice, which is reversed by *L. plantarum* D266.^39^ However, there is no exact molecular mechanism about how the probiotics influence the alteration of the ENS. Few studies have evaluated the role of enteric cholinergic signaling in probiotics-mediated colitis remission. With research advances as described above, further investigation is need to determine whether neuroimmune pathway may considered as a new target for probiotic-based therapies in regulating gastrointestinal disorders.

In this study, we discovered the abundance of *Lactobacillus* is negatively correlated with the disease phenotypes of UC patients and colitis mice. By screening *Lactobacillus* isolates from fermented foods both in vitro and in vivo experiments, we found that *L. plantarum* A736 effectively rescued colitis-related phenotypes mainly by increasing the synthesis of tryptophan-derived metabolite indole-3-lactic acid (ILA). We further confirmed the production of ILA released by *L. plantarum* A736. Moreover, *L. plantarum* A736 and ILA modulated intestinal immune via choline acetyltransferase-expressing (ChAT^+^) neurons in enteric nervous system (ENS), which exert protective effects on colitis symptoms. Overall, our findings point to the interplay between ENS and probiotics, and attempt to reveal the probiotics for colitis prevention from a new perspective.

## STAR METHODS

### RESOURCE AVAILABILITY

#### Lead contact

Further information and requests for resources and reagents should be directed to and will be fulfilled by the lead contact, Xuebo Liu.

#### Materials availability

Microbial strains used in this study are available from the lead contact upon request.

### EXPERIMENTAL MODEL AND STUDY PARTICIPANT DETAILS

#### Human participants

Fecal samples, endoscopic images and blood of 26 healthy controls (HC) and 35 ulcerative colitis (UC) patients at baseline (that is, before the screening colonoscopy) according to the inclusion and exclusion criteria described in **Table S1**, were obtained from Shaanxi Provincial People’s Hospital after written informed consent, which was approved by the Medical Ethical Committee of Shaanxi Provincial People’s Hospital (2023R109). The summarized clinical characteristics of all participants are listed in **Table S2**.

#### Animals

Six-to eight-week-old male C57BL/6 mice were obtained from SPF Biotechnology Co., Ltd (Beijing, China) and allowed to acclimatize to the animal facility environment for 2 weeks before experiment. ChAT-Cre mice and tdTomato^fl/stop/fl^ mice were purchased from GemPharmatech Co., Ltd (Jiangsu, China). ChAT-tdTomato mice were generated by crossing ChAT-Cre mice with a cell lineage reporter mouse line tdTomato^fl/stop/fl^ mice. All mice were maintained in individually ventilated cages (IVCs) under specific pathogen-free conditions. Protocols were approved by the animal ethics committee of Northwest A&F University (Ethic No XN2024-0414).

#### Bacterial strains and culture

All *Lactobacillus* spp. strains used in this study were isolated from the fermented foods, which are listed in **Table S3**. The strains from glycerol tubes were grown in Man Rogosa Sharpe agar (MRS) and then anaerobically cultured at 37 □ for 72 h, after which visible colonies were selected and transferred into MRS broth for further cultivation at 37 □ under anaerobic conditions.^40,41^ To prepare high-temperature inactivated *L. plantarum* A736, the bacteria were was heat-killed at 121□ for for 2 h.^42^

For bacterial supernatants experiments, the cultured *L. plantarum* A736 solution was centrifuged at 6000xg for 10 min in 4 °C to get supernatants and then followed by filtration through a sterilized 0.22-µm filter to remove residual bacterial cells.^43^ To interrogate the ability of strain to catabolize dietary tryptophan into indole derivatives, *L. plantarum* A736 were grown in MRS broth overnight, harvested by centrifugation, washed with PBS, and resuspended in sterile 10 mL of peptone-tryptone water (10 g/L peptone and 10 g/L tryptone, 5 g/L NaCl) supplemented with 0.6 mM L-tryptophan, then anaerobically incubated at 37 °C for 14 h. Bacteria cultures were centrifuged at 5000xg for 10 min, supernatant collected and filter by a sterilized 0.2 _μ_m pore diameter cellulose acetate filter (VWR) to further analysis.^44,45^

#### Cell cultures

The human colon carcinoma Caco-2 cell line were cultured in Dulbecco’s modified Eagle’s medium (DMEM) containing 10% (v/v) fetal bovine serum, 1% (v/v) penicillin/streptomycin solution, 4 mM L-glutamine and 1 mM sodium pyruvate, which were cultivated at 37°C with 5% CO_2_.^46,47^ For cultivation of Caco-2 cell monolayers, approximately 300000 cells were seeded for each 12 mm-diameter polycarbonate membrane insert and cultured for 21 days until the the transepithelial electrical resistance (TEER) had stabilized.^48,49^

### METHOD DETAILS

#### DSS-induced colitis model and treatments

The acute colitis was induced by dextran sodium sulfate (DSS) on the basis of previous procedures,^50^ which could better reflect the nature of IBD in human, especially resembling some features of flares UC. To induce colitis, mice received 2.5 % DSS in drinking water from day 0 until day 7. DSS was made fresh every other day throughout the experiment. All mice were daily weighed and assessed for stool consistency and rectal bleeding, which could be used to evaluate a disease activity index (DAI). In addition, colitis severity without sacrificing the mice was performed by colonoscopy and in vivo imaging after the 7 days of DSS induction (day 7 or 8). For bacterial treatment, mice were orally administered with fourteen *Lactobacillus* strains (*L. plantarum* E853, *L. plantarum* D333, *L. plantarum* A736, *L. paracasei* A7411, *L. paracasei* C941, *L. paracasei* E857, *L. fermentum* E451, *L. fermentum* E464, *L. brevis* B651, *L. brevis* C231, *L. delbrueckii* O171, *L. delbrueckii* E1705, *L. casei* J3704, *L. casei* C5603) at a sufficient dose (11710^9 CFU/0.2 mL) per day for 1 weeks, or sterilized PBS as a DSS group or a single dose of 5-ASA (100 mg/kg) as a positive drug control group, then subjected to DSS for model induction, and continuously given probiotic until mice were sacrificed. Moreover, mice were treated only sterilized PBS without DSS induction as a control group. For the protective component of *L. plantarum* A736, mice were treated with *L. plantarum* A736, heat-killed *L. plantarum* A736, the supernatant of *L. plantarum* A736 and MRS medium for 1 week, followed by DSS exposure. For ILA treatment experiments, mice were gavaged with 40 mg/kg ILA or PBS daily for one week and simultaneously induced by DSS.^51^ For dietary studies, mice were fed with either low tryptophan (0.021%) or high tryptophan diet (1.47%) 2 weeks prior to DSS induction and maintained on their respective diets continuously until the end of experiment.^52^ For non-dietary studies, mice received standard chow containing 0.21% tryptophan based on American Institute of Nutrition (AIN)-93 recommendations.^53^ Composition of diets was demonstrated in **Table S4**. For the ablation of enteric ChAT-expressing neurons, mice were injected with the virus and treated *L. plantarum* A736 or ILA three weeks later, followed by DSS exposure.

#### AB/PAS staining analysis

Colon tissues containing fecal pellets were collected from mice, fixed with methanol-Carnoy’s fixative (60% methanol, 30% chloroform, 10% glacial acetic acid) at least 24 hours and then processed into paraffin-embedded sections.^54^ Tissues sections were stained with Alcian blue and Periodic acid-Schiff (AB/PAS) to evaluate goblet cells.

#### Intra-colonic injection of AAV viruses

Mice were anesthetized and ophthalmic ointment was placed over the eyes to prevent dehydration. After shaving and sterilization of the abdominal area, the mice were positioned on a stereoscope and covered with a sterile surgical drape. The colon was exteriorized by a midline incision through the abdominal wall. 2μL of AAV virus (500 nL*4 sites) was injected with a pulled glass pipette, and the needle tip was left in place for at least 5 minutes to prevent reflux.^55–58^ Following the procedure, the abdominal wall was closed using absorbable sutures and then the skin was closed as above. Three weeks after virus injections, we further assessed the effect of *L. plantarum* A736 or ILA to DSS induction.

#### Growth conditions of strains

For growth curve assay, the *Lactobacillus* strains were inoculated in sterile microplate containing MRS medium and maintained at 37 °C. Absorbance values were recorded at 600nm (OD_600 nm_) with an interval of 1 h over 24 h for all bacterial strains by using Bioscreen C.

#### Simulated gastrointestinal tolerance test

For gastrointestinal tolerance experiments, the *Lactobacillus* strains were exposed sequentially to the simulated gastrointestinal fluid (3 h) and simulated intestinal fluid (4 h) as previously reported.^59^ Then the survival of *Lactobacillus* strains was monitored by plate counting at 0 h and 7 h (3 h plus 4 h).

#### Antimicrobial activity

For antibacterial activity of the *Lactobacillus* strains against pathogen bacteria such as *Staphylococcus aureus* ATCC 6538 and *Escherichia coli* ATCC 700928, the Oxford cup method was performed according to previous study.^39,46^ Petri plates were poured with sterile Luria Bertani (LB) agar medium and seeded with 200 µL freshly grown pathogen bacteria, followed by wells of 9 mm diameter excavated with a sterilized Oxford cup. Then 200 µL *Lactobacillus* cultures grown to a stationary phase or MRS broth solution (as the blank control) were loaded per well. After plates were incubated at 37 °C for 24 h, a vernier caliper was used to evaluate the diameter of inhibition zones.

#### Adhesion of strains

For bacterial adhesion experiments, *Lactobacillus* strains were washed three times with PBS and resuspended in DMEM containing 10% fetal bovine serum, which were added to polarized Caco-2 monolayers at a multiplicity of exposure (MOE) of ∼10 for strains. Following 1 h of incubation, monolayers were rinsed in DMEM, lysed with 0.1% Triton X-100 in PBS and adherent bacteria were enumerated by plating serial dilutions on Lactobacillus selective MRS agar plates to count as previously described.^46^

#### Paracellular permeability assay

For analysis of epithelial permeability, Caco-2 cell monolayer pretreated with *Lactobacillus* strains (MOE; 10, 24 h) before TNFα exposure (20 ng/mL, 24 h) was supplemented in the basolateral chamber to mimic colitis situation in vitro as previous protocol.^46,60^ Subsequently, 4 kDa FITC-dextran was added to the apical side at 1 mg/ml in HBSS and translocation of FD4 to the basolateral compartment was measured after 2 h incubation using a Spark Multimode Microplate Reader (Tecan Austria GmbH, Austria).

#### Metabolomic analysis

For targeted metabolomics of neurotransmitter, colonic tissue was homogenized in methanol solutions. Following centrifugation and filtration, the sample extract was analyzed by LC-MS/MS using ACQUITY UPLC HSS T3 C18 column. The analytical conditions follow: solvent A was water with 0.1% formic acid, and solvent B was acetonitrile with 0.1% formic acid. The gradient conditions were as follows: 0 min, 5% B; 0 to 8 min, gradient to 95% B; 8 to 9.5 min, 95% B; 9.6 to 12 min, gradient to 5% B. The flow rate was 0.30 ml/min.

For non-targeted metabolomics assay, feces and *L. plantarum* A736 supernatant was homogenized in the methanol buffer. The resulting supernatant dried, redissolved and filtered before LC-MS detection. Metabolites were measured on the UltiMate 3000 UPLC (Thermo Fisher Scientific, Bremen, Germany) coupled with a high-resolution tandem mass spectrometer Q-Exactive (Thermo Fisher Scientific, Bremen, Germany), which equipped with an ACQUITY UPLC T3 column. The analytical conditions follow:solvent A was water plus ammonium acetate and acetic acid, and solvent B was acetonitrile. The gradient conditions were as follows: 0 to 0.8 min, 2% B; 0.8 to 2.8 min, gradient to 70% B; 2.8 to 5.6 min, 70% to 90% B; 5.6 to 6.4 min, 90% to 100% B; 6.4 to 8.0 min, 100% B; 8.0 to 8.1 min, 100% to 2% B; 8.1 to 10 min, gradient to 2% B. The flow rate was 0.30 ml/min. Data collection was operated in both positive and negative ion modes within a range of 70-1050 m/z. Metabolic identification was validated by searching in-house standard database and integrated public databases.

For targeted metabolomics of the indole derivatives, feces and *L.* pl*antarum* A736 culture supernatant underwent homogenization in methanol solutions before collecting the supernatant, which was then dried down. Following resuspension and filtration, the samples were analyzed by LC-MS using a C18 column. The analytical conditions follow: solvent A was water and solvent B was acetonitrile. The gradient conditions were as follows: 0 to 3 min, 5 % to 30 % B; 3 to 7 min, gradient to 100 % B; 7 to 8 min, 100% B; 8 to 9 min, gradient to 5 % B; 9 to 10 min, 5 % B. The flow rate was 0.30 ml/min.

#### Isolation of colonic lamina propria leukocytes

For isolation of colonic lamina propria immune cells, the colon was opened longitudinally and cut into small pieces. The tissue segments pieces were incubated with dissociation buffer containing fetal bovine serum, dithiothreitol (DTT) and EDTA at 37°C for 20 min under continuous rotation, which aimed to remove epithelial cells, intraepithelial lymphocytes and mucus. Then, the remaining pieces were washed thoroughly in cold PBS and digested in digestive solution with fetal bovine serum, DNase I and Collagenase for 45 min at 37°C shaking. The digested cell suspension was passed sequentially through 100 μm cell strainers and resuspended in 40% Percoll, followed by overlaying on top of the 80% Percoll solution. After centrifugation, colonic lamina propria immune cells were obtained from the interface of 40/80% Percoll solution.^14,22,61^

#### Flow cytometry analysis

Immune cells were isolated from colonic lamina propria as described above. For cell-surface immunostaining, single-cell suspensions were stained with the following antibodies against CD45, CD11b, Ly-6C, F4/80, Ly-6G, CD11c, MHC□, CD3ε, CD4, CD25 (see antibody table for details). For intracellular staining, after cell surface staining, cell suspensions were fixed, permeabilized using transcription factor buffer sets according to the manufacturer’s protocol and stained with antibodies including anti-RORγt and anti-FOXP3. The flow cytometry data were collected with a BD FACSAria™ III and analyzed with FlowJo software.

#### Isolation of myenteric and submucosal plexus and immunofluorescence

Freshly dissected colons were gently cleaned with ice-cold PBS and cut open longitudinally. Separation of myenteric and submucosal plexus was dependent on previous report.^16,62^ The myenteric and submucosal plexus was then pinned to Sylgard-coated plate with Minutien pins, fixed for 16 h using 4% paraformaldehyde (PFA) at 4 □. After washing in PBS, tissues were filled with permeabilization buffer (0.2% Triton-X in PBS) for 2 h, blocking buffer (0.2% Triton-X, 5% bovine serum albumin and 5% donkey serum in PBS) for 1 h on an orbital shaker at room temperature, and incubated with primary antibodies including anti-HuC/D (1:500; ab184267; Abcam) and anti-ChAT (1:100; AB144P; Sigma-Aldrich) in the same blocking buffer for 48 h at 4 □. Next, tissue was incubated with secondary antibodies using the following antibodies: Donkey anti-Rabbit (1:200; A32790; Thermo Fisher Scientific) and Donkey anti-Goat (1:200; A32816; Thermo Fisher Scientific). Imaging was obtained using a LEICA TCS SP8 confocal microscope (Leica Microsystems, Germany) with a 63× oil objective, 1 zoom, 1 airy unit pinhole, and 1024×1024 pixels xy resolution. The number of neurons per sample was subsequently quantified. For each sample, 5 ganglia were randomly selected and quantified. Data for each mouse is composed of the mean of the 5 ganglia counted.

#### RNA sequencing analysis

The total RNA from colonic tissues was extracted with TRIzol reagent. The quality and concentration of the RNA was determined by using the Bioanalyzer 2100 (Agilent Technologies, CA, USA). After library preparation, RNA sequencing was performed with Illumina NovaSeq 6000 platform by Novogene Company (Beijing, China). TrimGalore (version 0.6.6) was used to remove the adapter and filter out low-quality reads. STAR (version 2.7.10a) was used to align the clean reads to the mouse reference genome mm39, and StringTie (version 2.1.6) was used for quantification to obtain the gene count matrix. Genes fulfilling the criteria p < 0.05 and log2 fold change > 1 were considered to be differentially expressed using the Limma R package. Differentially expressed genes (DEG) were utilized as the inputs for Gene Set Enrichment Analysis (GSEA) and KEGG pathway analysis by ClusterProfiler R package (version 4.10.1).

#### Whole**□**genome sequencing

Genomic DNA was extracted from *L. plantarum* A736 using bacterial DNA extraction kit (Majorbio, shanghai, China). For Illumina sequencing libraries, genomic DNA was fragment at 400-500 bp. For PacBio sequencing libraries, DNA samples were sheared into ∼10kb fragments. Then genomic DNA was sequenced using a combination of PacBio and Illumina sequencing platforms by Shanghai Majorbio Bio-pharm Biotechnology Co., Ltd. (Shanghai, China). Next, the clean short reads and HiFi reads were assembled to construct complete genomes using Unicycle v0.4.8, followed by Pilon v1.22 to polish the assembly using short-read alignments. The coding sequences (CDs) were predicted using Prodigal and GeneMarkS. Also, the predicted CDs were annotated from KEGG database.

#### 16S rRNA sequencing

Total fecal DNA was extracted from mice and humans, followed by PCR amplification of the bacterial 16S rRNA gene targeting the variable regions 3 and 4 (V3–V4) with primer pairs: 338F (5’-CCTACGGGNGGCWGCAG-3’) and 806R (5’-GACTACHVGGGTATCTAATCC-3’). Sequences were generated by an Illumina NovaSeq platform at LC-Bio Technology Co. Ltd. (Hangzhou, China). The raw sequences were processed and clustered into amplicon sequence variants (ASVs) using DADA2. The sequencing data were analyzed mainly by QIIME2 and lianchuan omics. Afterward, the taxonomy of these features was assigned to the NCBI database with a 70 % confidence threshold.

#### Quantitative RT-PCR

Total RNA from mouse colon was extracted according to the RNAex Pro Reagent (Accurate Biology, AG21102), followed by converting into cDNA using a reverse transcription kit (Accurate Biology, AG11728). Subsequently, qRT-PCR was performed following the instructions provided with the SYBR Green Premix Pro Taq HS qPCR Kit (Accurate Biology, AG11701). Specific primer sequences can be found in Key Resource Table. The PCR program was set under the following conditions: 95 □ for 30 sec, followed by 40 cycles of 95 □ for 5 sec and 60 □ for 30 sec. The relative mRNA expression level was calculated using the 2-ΔΔct method by selecting β-actin as the internal reference gene.

#### In vivo imaging

Referring to published studies, intestinal inflammation may optionally be noninvasively measured by in vivo imaging using the luminol-based chemiluminescent probe L-012.During anesthesia, 25 mg/kg of L-012 solution was administered intraperitoneally.^63,64^ Next, processed mice were placed into the the imaging chamber of the IVIS Spectrum system (PerkinElmer, America) to acquire images. For quantitative analyses, the luminescent signal in the standardized regions of interest (ROI) was calculated by the Living Image software.

#### Histology and immunofluorescence

Colon samples was fixed in 4% paraformaldehyde, embedded in paraffin and sectioned. For histopathological evaluation, sections were stained with hematoxylin and eosin (H&E) followed by scoring according to the published standard.^50^ For immunostaining, slides were blocked, permeabilized and then incubated in primary antibodies (rabbit anti-Occludin, Abcam, ab216327, 1:200; mouse anti-Claudin-1, Santa Cruz, sc-166338, 1:500; mouse anti-Mucin 2, Santa Cruz, sc-7314, 1:500; rat anti-ZO-1, Santa Cruz, sc-33725, 1:500) as published method.^65^ Primary antibodies were visualized using secondary fluorescently labelled antibodies as shown in the Key Resource Table. After washing, the nucleus was counterstained with DAPI and images were obtained. Mean fluorescence intensity was quantitated by Image J software.

### QUANTIFICATION AND STATISTICAL ANALYSIS

Data are expressed as the mean ± standard error of mean (SEM) from at least three independent biological replicates. Representative immunostaining, histology and flow cytometry images are shown. Differences between groups was determined by unpaired Student’s t-test and one-way ANOVA with Tukey’s multiple comparison test by GraphPad Prism. P < 0.05 was considered statistically significant.

#### Results

##### Lactobacillus abundance negatively correlates with clinical features of UC

To evaluate the connection of the gut microbiome and UC development in human, we collected and analyzed blood and stool samples from UC patients (n=35) and healthy controls (HC, n=35) previously recruited by us **(Table S1 and Figure 1A**). Endoscopically, patients with UC feature loss of vascular pattern, friability of the mucosa, erosions, and, in the context of severe inflammation, spontaneous bleeding and ulcerations (**Figure 1B**). For disease activity parameters, there were an elevation in C-reactive protein (CRP) in patients (HC<0.5 mg/L), whereas diminishing the levels of hemoglobin **(Table S2 and Figure 1C**). In addition, we also observed increased leukocyte counts, including monocytes and neutrophil in patients (**Figure 1D-1F**). Collectively, these results show classic clinical characteristics in UC patients.

**Figure 1.**
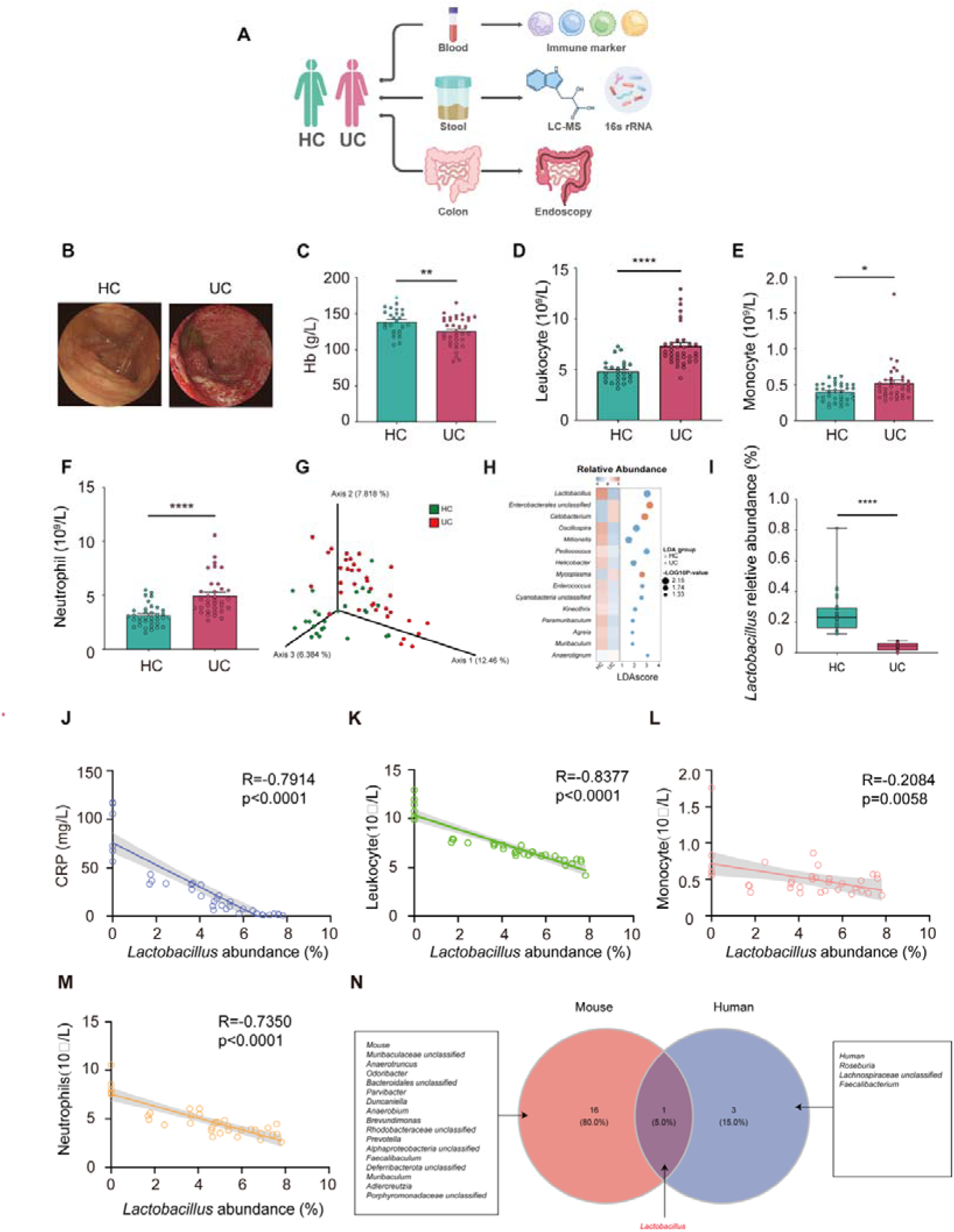
*Lactobacillus* abundance negatively correlates with UC progression in humans. (A) Schematic illustration from human subjects. Ulcerative colitis patients (UC, n=35), healthy controls (HC, n=26). (B) Representative endoscopic images. (C) Levels of hemoglobin in the blood. (D-F) Leukocyte (D), monocyte (E) and neutrophil (F) counts in the blood. (G) Principal co-ordinates analysis (PCoA) of unweighted unifrac distance based on the 16S rRNA sequencing from stool samples. ANOSIM (analysis of similarities), R = 0.1, p = 0.007. (H) Linear discriminative analysis (LDA) score of the top 15 most differential genera from HC and UC. (I) Relative abundance of *Lactobacillus* genus from HC and UC. (J-M) Correlative analysis of *Lactobacillus* abundance with C-reactive protein (CRP) values (J), leukocyte (K), monocyte (L) and neutrophil (M) counts in UC patients. (N)Venn diagram indicating genus enriched in the stool samples from healthy people and normal mice (LDA score > 3 and p < 0.05 using Wilcoxon rank-sum test). Mean ± SEM shown. ns, not significant, *p < 0.05, **p < 0.01, ***p < 0.001, and ****p < 0.0001, determined by unpaired Student’s t-test. See also Figure S1, Table S1 and S2.

We next conducted 16S ribosomal RNA (rRNA) sequencing on stool samples from both HC and UC patients followed by profiling of gut microbial communities. As previous report, beta diversity represented by principal coordinates analysis (PCoA) exhibited significant clustering separation among the two groups (**Figure 1G**). Furthermore, *Lactobacillus* displayed a most differential abundance from groups and reached the LDA score of 3.03 by the Linear discriminant analysis of effect size (LEfSe) analysis (**Figure 1H**). Differentiated-abundance analysis showed that UC patients markedly reduced the relative abundance of *Lactobacillus* genus, which is consistent with a recent report (**Figure 1I**).^66^ We further explored the relationship between *Lactobacillus* and clinical features of UC by performing Pearson correlation analyses. Remarkably, a negative correlation was observed for the abundance of *Lactobacillus* and several markers of disease activity, including inflammatory markers CRP, and inflammatory cell counts in the blood (**Figure 1J-1M**). Consistent with these findings, then abundance of *Lactobacillus was* negatively associated with disease course by assessing a clinical cohort based on UC patients **(Figure S1A)**.^67^ Such a phenomenon is also well aligned with the results from our animal studies on gut microbial signatures in dextran sulfate sodium (DSS)-induced colitis mice. All colitis mice exhibited the lower abundance of *Lactobacillus compared with the control mice*, and the abundance of *Lactobacillus* negatively correlated with disease-related indicators **(Figure S1B-S1K)**. Specifically, although most elevated bacterial genus of the HC differed from Control mice (LDA score > 3 and p < 0.05 using Wilcoxon rank-sum test), an increase in the *Lactobacillus* genus in HC was same as Control mice (**Figure 1N**). Taken together, these clinical and animal results provide evidence for the link between *Lactobacillus* abundance and UC progression, further implying the possibility that the deficiency of *Lactobacillus* genus may contribute to colitis development.

##### L. plantarum A736 mitigates acute colitis and regulates intestinal immune in mice

*Lactobacillus* is a genus of beneficial bacteria commonly found in fermented foods. To identify promising *Lactobacillus* strains with potential for colitis remission, we isolated 50 strains from 14 different *Lactobacillus* species derived from fermented foods in China (**Table S3**). We first explored the fundamental probiotic properties of the strains and their ability to regulate intestinal barrier function under *in vitro* conditions. Through simulated gastrointestinal tract analysis, antimicrobial activity assessment, bacterial adhesion evaluation, and epithelial permeability profiling in polarized Caco-2 monolayers, we observed strain-specific differences in probiotic characteristics and intestinal barrier function (**Figure S2A–S2S**). Using principal component analysis (PCA) and fuzzy synthetic evaluation (FSE) (**Figure S2T–S2V**), we identified 14 *Lactobacillus* strains with favorable probiotic properties.

We next attempted to identify whether 14 *Lactobacillus* strains could attenuate colitis *in vivo*. To this end, we modeled colitis in mice induced by DSS, as with clinical features of UC patients. Notably, oral administration of *L. plantarum* A736, *L. paracasei* A7411, or *L. delbrueckii* E1705 significantly attenuated weight loss, reduced the disease activity index, improved histological scores, and increased colon length **(Figure S3A-S3I and Figure 2A**). Additionally, we found that all three strains significantly suppressed the mRNA expression of intestinal proinflammatory cytokines, including TNF-α and IL-1β, which were elevated in DSS-induced colitis mice **(Figure S3J-S3K)**. To evaluate gut mucosal barrier integrity, including the mucus barrier and epithelial barrier, we quantified the number of mucus-producing goblet cells and assessed the expression of Muc2 and tight junction proteins (Claudin-1, ZO-1, and Occludin) in colonic tissues. Compared to colitis mice, treatment with *L. plantarum* A736, *L.* A7411 or *L. delbrueckii* E1705 resulted in a significant upregulation of these markers **(Figure S3L-S3P and Figure S4A-S4I)**. Based on PCA and FSE analysis, we observed that *L. plantarum* A736 exhibited the greatest potential for alleviating colitis **(Figure S4J-S4K)**. Remarkably, we found that the colitis severity was reduced in the *L. plantarum* A736 colonized mice by performing endoscopy and in vivo imaging using the chemiluminescent probe L-012 (**Figure 2B-2D**). *L*. *plantarum* A736 demonstrated effective colonization in the intestine of DSS-induced colitis mice, as confirmed by 16S rRNA sequencing analysis following oral gavage (**Figure 2E**). Given that UC is characterized by chronic immune-mediated inflammatory condition of the large intestine,^68,69^ we analyzed the immune cell subpopulations in colonic lamina propria (cLP) by flow cytometry **(Figure S5A-S5C)**. *L. plantarum* A736 supplementation significantly reduced the percentage of monocytes, macrophages, neutrophils, Th17 cells relative to DSS exposure. However, no significant alterations were shown in the percentage of both dendritic cells and Treg cells, supporting that *L. plantarum* A736 contributed to the balance of Th17/Treg response (**Figure 2F-2L**). Altogether, these data strongly indicate that administration of *L. plantarum* A736 improves the phenotype of colitis, colonic mucus barrier, gut epithelium and intestinal immune. Consequently, we further focused on *L.plantarum* A736 to investigate the underlying mechanisms.

**Figure 2.**
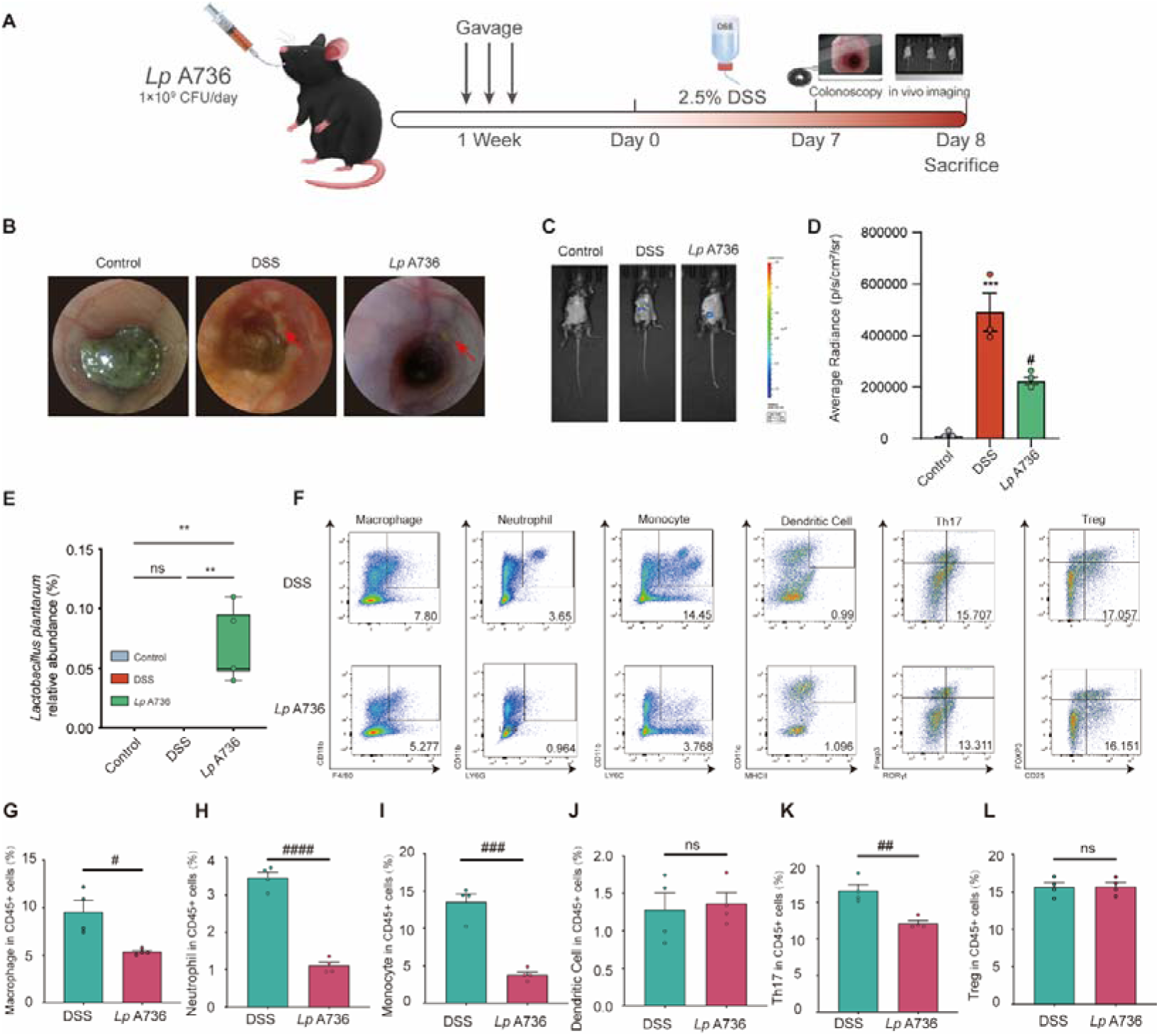
*L. plantarum* A736 protects mice against DSS-induced colitis and regulates intestinal immune. (A) Schematic illustration of *L. plantarum* A736 treatment paradigm. (B) Representative colonoscopy images (n=3). (C-D) Representative image captured by an in vivo imaging system and quantification (n=3). (E) Relative abundance of *L. plantarum*. (F) Representative flow cytometry plots for monocytes, macrophages, neutrophils, dendritic cells, Th17 and Treg cells from colonic lamina propria (cLP). (G-L) The percentage of monocytes, macrophages, neutrophils, dendritic cells, Th17 and Treg cells (n=4). Control group (Control), DSS group (DSS) and *L. plantarum* A736 treatment group (*Lp* A736). Mean ± SEM shown. Data are determined by one-way ANOVA with Tukey’s multiple comparison test (D) unpaired Student’s t-test (G-L) or by Kruskal-Wallis test (E). ns, not significant, *p < 0.05, **p < 0.01, ***p < 0.001, and ****p < 0.0001, compared with the Control group, ^#^p<0.05, ^##^p<0.01, ^###^p < 0.001, and ^####^p < 0.0001, compared with the DSS group. See also Figures S2, S3 and S4, Table S3.

##### L. plantarum A736 regulates ChAT-expressing neurons

To elucidate the mechanisms underlying the protective effects of *L. plantarum* A736 on colitis symptoms, we conducted RNA sequencing on colon tissues. Analysis revealed that DSS exposure resulted in the upregulation of 1765 genes and the downregulation of 1322 genes compared to the control group **(Figure S6A)**. Kyoto Encyclopedia of Genes and Genomes (KEGG) and Gene Set Enrichment Analysis (GSEA) further demonstrated that the neuroactive ligand-receptor interaction pathway was significantly suppressed by DSS induction **(Figure S6B-S6C)**. Strikingly, in contrast to DSS induction, *L. plantarum* A736 administration led to the upregulation of 1344 genes and the downregulation of 574 genes (**Figure 3A**). KEGG pathway analysis revealed that the differentially expressed genes (DEGs) were predominantly enriched in the neuroactive ligand-receptor interaction pathway, which emerged as the top-ranked pathway (**Figure 3B**). Additionally, GSEA confirmed the activation of the neuroactive ligand-receptor interaction pathway following *L. plantarum* A736 treatment (**Figure 3C**). These findings underscore the critical role of neuroactive signaling pathways in mediating the protective effects of *L. plantarum* A736 against colitis.

**Figure 3.**
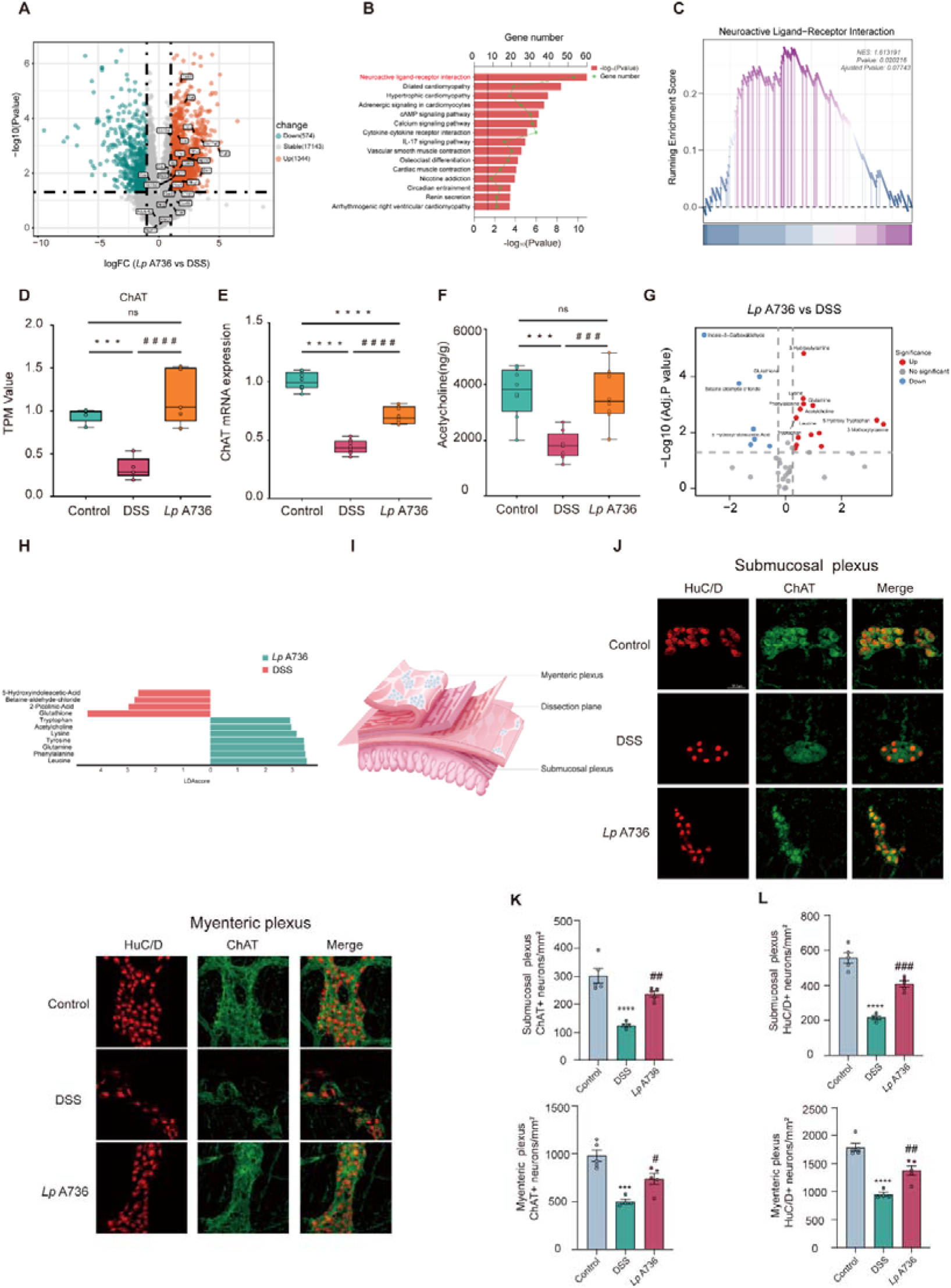
*L. plantarum* A736 impacts neuroactive signaling pathways and neurotransmitter production. (A) Volcano plot displaying differentially expressed genes in colon tissues between *Lp* A736 and DSS (n=8). (B) KEGG analysis demonstrating distinct differences in pathways between *Lp* A736 and DSS (n=8). (C) GSEA analysis of neuroactive ligand-receptor interaction pathway□associated genes in *Lp* A736-treated mice and DSS-induced mice (n=8). (D) Transcripts per kilobase million (TPM) values for ChAT by RNA sequencing (n=8). (E) The mRNA expression of ChAT in colon tissues accessed by quantitative real-time PCR (n=8). (F) The level of ACh in colon tissues (n=8). (G) Volcano plot showing differential neurotransmitters in colon tissues between *Lp* A736 and DSS (n=8). (H) Linear discriminative analysis (LDA) score of differentially enriched neurotransmitters from LEfSe analysis between *Lp* A736 and DSS (n=8). (I) Experimental diagram for isolation of submucosal and myenteric plexus. (J) ChAT (green) and HuC/D (red) immunostaining of colonic submucosal and myenteric plexus (n=4-5). Scale bar, 36.8 μm. (K-L) Quantification of ChAT^+^ and HuC/D^+^ neurons in both submucosal (K) and myenteric plexus (L) of the colon (n=4-5). Control group (Control), DSS group (DSS) and *L. plantarum* A736 treatment group (*Lp* A736). Mean ± SEM shown. Data are determined by one-way ANOVA with Tukey’s multiple comparison test(K-L). ns, not significant, *p < 0.05, **p < 0.01, ***p < 0.001, and ****p < 0.0001, compared with the Control group, ^#^p<0.05, ^##^p<0.01, ^###^p < 0.001, and ^####^p < 0.0001, compared with the DSS group. See also Figures S6.

Emerging evidences have shown that enteric neurons play key roles in mediating gastrointestinal diseases.^26^ Enteric neurons comprise distinct neuron subtypes that have different functions and work cooperatively to regulate most aspects of intestinal physiology.^70^ We further profiled distinct enteric neuron subsets from mice using previous single-cell RNA sequencing data,^12,71^ which revealed that cholinergic neurons (termed ChAT-expressing neurons) were considered as major neuron subtypes of enteric neurons based on the expression of specific marker genes **(Figure S6D-S6G)**, which is consistent with the previously reported.^16^ Remarkably, our transcriptome analysis demonstrated the higher expression of ChAT in colonic tissues from *L. plantarum* A736-adiministered mice relative to DSS-induced mice, which were also verified by quantitative real-time PCR assays (**Figure 3D-3E**). We next profiled neurotransmitter in the colon tissue through targeted metabolomic data. Acetylcholine (ACh), the primary neurotransmitter produced by cholinergic neurons, was identified as one of the top-ranked metabolites significantly enriched in the *L. plantarum* A736-treated group in comparison to DSS group (**Figure 3G-H, Figure S6H-6I)**. Additionally, oral gavage of *L. plantarum* A736 dramatically improved the level of ACh relative to DSS administration (**Figure 3F**). These findings demonstrate that *L. plantarum* A736 ameliorates colitis through a cholinergic neuron-dependent mechanism.

To further investigate the effects of *L. plantarum* A736 on neuronal populations, we generated ChAT-Cre;tdTomato^fl/stop/fl^ reporter mice to visualize gut-innervating cholinergic neurons. Immunofluorescence analysis of colon sections revealed ChAT-tdTomato^+^ neurons (co-localized with neuronal marker βIII-tubulin) in close proximity to EpCam1^+^ epithelial cells within crypts and muscularis layers, both at steady state and during intestinal inflammation **(Figure S6J-L)**. Quantitative whole-mount analysis of submucosal and myenteric plexuses demonstrated significant depletion of ChAT+ neurons and total enteric neurons (HuC/D^+^) in DSS-treated mice. Strikingly, *L. plantarum* A736 administration not only restored ChAT^+^ neuron numbers but also increased overall enteric neuron density (**Figure 3I-L and Figure S6M)**. These findings uncover a novel neuroprotective mechanism of probiotic action, establishing cholinergic neurons as crucial mediators of *L. plantarum* A736’ s therapeutic effects in colitis.

##### *L. plantarum* A736 regulates ChAT-expressing neurons via tryptophan metabolism

To better identify the protective component of *L. plantarum* A736, we administered one of four treatments prior to DSS exposure: (1) live *L. plantarum* A736 (*Lp* A736), (2) heat-inactivated *L. plantarum* A736 (HK-*Lp* A736), (3) *L. plantarum* A736 culture supernatant (*Lp* A736-sup), or (4) the standard culture medium (Man-Rogosa-Sharpe, MRS) as control (**Figure 4A**). We found only live *L. plantarum* A736 and *L. plantarum* A736 supernatant treatment had a comparable effect to improve colitis phenotypes, restored gut mucosal barrier (Claudin-1, ZO1, Occludin and Muc2), downregulated pro-inflammatory cytokines expression (TNF-α, IL-17, IL-6 and IL-1β) (**Figure 4B-4I**). These results indicate that the anti-colitic effects were mediated primarily by *L. plantarum* A736-derived metabolites rather than by direct bacterial components.

**Figure 4.**
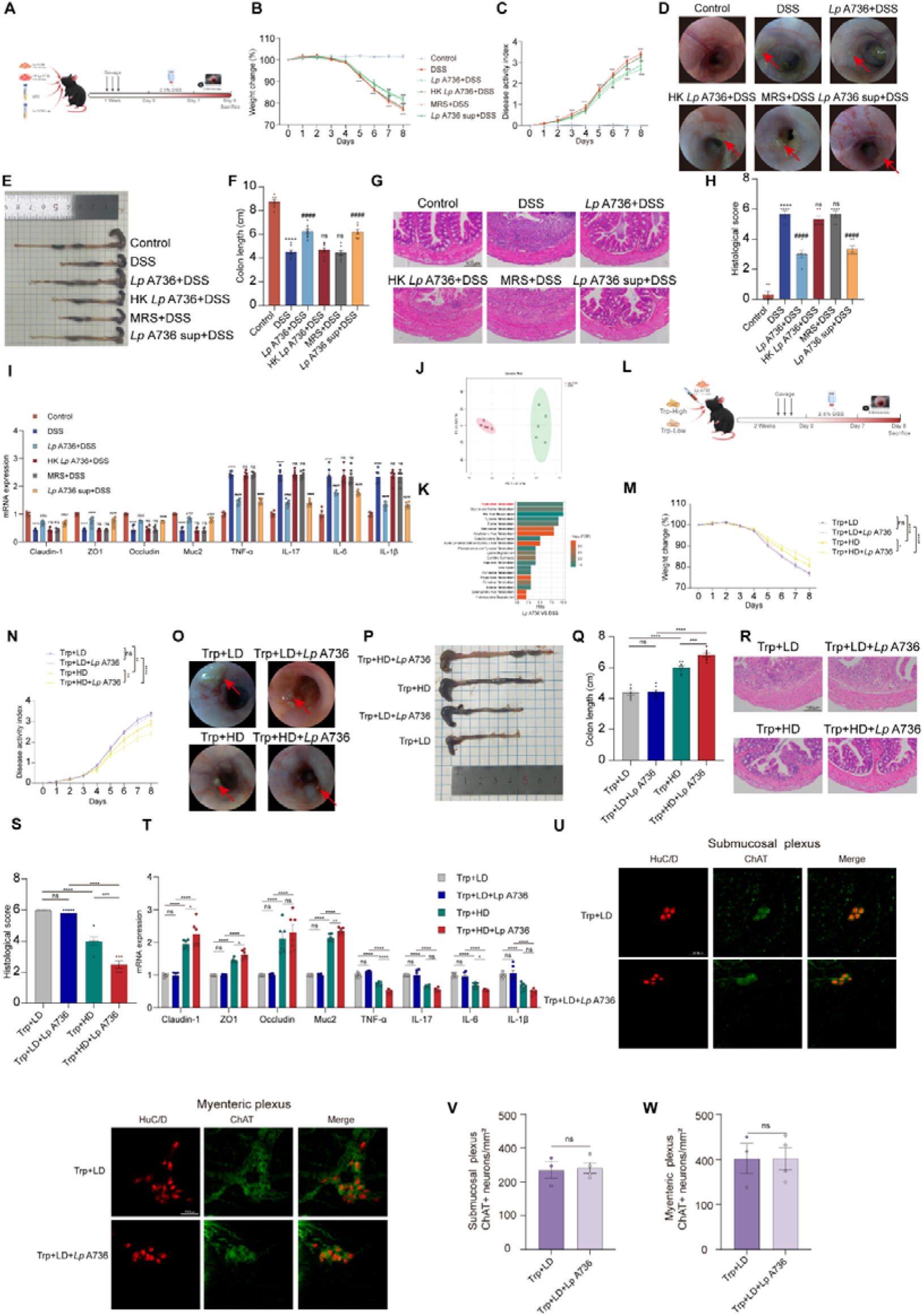
*L. plantarum* A736 alleviates colitis and regulates ChAT-expressing neurons via tryptophan metabolism. (A) Experimental scheme depicting mice orally administered live *L. plantarum* A736 (*Lp* A736), or heat-killed *L. plantarum* A736 (HK *Lp* A736), or *L. plantarum* A736 supernatant (*Lp* A736 sup), or original culture medium (Man-Rogosa-Sharpe, MRS) or PBS. (B) Body weight change (n=8). (C) Disease activity index (DAI) score (n=8). (D) Representative colonoscopy images (n=3). (E) Representative photographs of colon. (F) Colon length (n=8). (G and H) Representative image of hematoxylin and eosin (H&E)-staining sections in colon and histological scores (n=6). Scale bar, 50 μm. (I) The mRNA expression of gut barrier-related proteins (Claudin-1, ZO1, Occludin and Muc2) and pro-inflammatory cytokines (TNF-α, IL-17, IL-6 and IL-1β) in colonic tissues evaluted by quantitative real-time PCR (n=6). (J) Principal component analysis (PCA) of stool samples by untargeted metabolome (n=5). (K) Pathway-associated quantitative enrichment analysis (QEA) of differentially enriched metabolites in stool samples between DSS and *Lp* A736 (n=5). (L) Diagram of tryptophan diet schedule in mice gavaged with *L. plantarum* A736 or PBS. (M) Body weight change (n=8). (N) Disease activity index (DAI) score (n=8). (O) Representative colonoscopy images (n=3). (P) Representative photographs of colon. (Q) Colon length (n=8). (R and S) Representative image of hematoxylin and eosin (H&E)-staining sections in colon and histological scores (n=6). Scale bar, 50 μm. (T) The mRNA expression of gut barrier-related proteins (Claudin-1, ZO1, Occludin and Muc2) and pro-inflammatory cytokines (TNF-α, IL-17, IL-6 and IL-1β) in colonic tissues evaluted by quantitative real-time PCR (n=6). (U) ChAT (green) and HuC/D (red) immunostaining of colonic submucosal and myenteric plexus (n=3-4). Scale bar, 36.8 μm. (V-W) Quantification of ChAT^+^ neurons in both submucosal (V) and myenteric plexus (W) of the colon (n=3-4). Control group (Control), DSS group (DSS), live *L. plantarum* A736 treatment group (*Lp* A736), heat-killed *L. plantarum* A736 (HK *Lp* A736), *L. plantarum* A736 supernatant (*Lp* A736 sup), original culture medium (Man-Rogosa-Sharpe, MRS). Low-tryptophan (Trp) diet (Trp LD), high-tryptophan diet (Trp HD), low-tryptophan diet and *L. plantarum* A736 (Trp LD+*Lp* A736), high-tryptophan diet and *L. plantarum* A736 (Trp HD+*Lp* A736). Mean ± SEM shown. Data are determined by two-way ANOVA with Tukey’s multiple comparison (B-C and M-N), one-way ANOVA with Tukey’s multiple comparison test (E-I and Q-T) or unpaired Student’s t-test (V-W). ns, not significant, *p < 0.05, **p < 0.01, ***p < 0.001, and ****p < 0.0001, compared with the Control group, ^#^p<0.05, ^##^p<0.01, ^###^p < 0.001, and ^####^p < 0.0001, compared with the DSS group. See also Figures S7.

To identify colitis-alleviating signals derived from *L. plantarum* A736, we performed untargeted metabolomic profiling of fecal samples. Comparative analysis revealed distinct metabolite profiles between DSS-treated and *L. plantarum* A736-supplemented mice (**Figure 4J, S7A)**. Quantitative enrichment analysis (QEA) identified tryptophan metabolism as the most significantly enriched pathway (**Figure 4K**). Furthermore, untargeted metabolomic analysis of the *L. plantarum* A736 culture supernatant revealed enrichment in tryptophan metabolism and differential production of metabolites, consistent with the findings observed in the fecal samples **(Figure S7B-S7D)**. Furthermore, whole-genome sequencing of *L. plantarum* A736 revealed significant enrichment of genes involved in amino acid metabolism and biosynthesis pathways **(Figure S7E-7F)**.

To determine whether tryptophan is required for *L. plantarum* A736-mediated colitis protection, mice were maintained on either high-or low-tryptophan diet beginning two weeks prior to DSS administration and continuing throughout the experiment (**Figure 4L**). Strikingly, the low-tryptophan diet completely abolished both the protective effects of *L. plantarum* A736 on colitis severity and its ability to preserve ChAT-expressing neurons (**Figure 4M-4W**). Moreover, we found that both high-tryptophan diet alone and its combination with *L. plantarum* A736 significantly improved colitis outcomes. Notably, the combined treatment exhibited additive protective effects, indicating that the high-tryptophan diet enhances L. plantarum A736-mediated protection against colitis. (**Figure 4M-4T**). Collectively, these results demonstrate that tryptophan metabolism is essential for the beneficial effect of *L. plantarum* A736 on both colitis progression and enteric neuron maintenance, highlighting a crucial functional connection between this microbial metabolic pathway and the enteric nervous system.

##### *L. plantarum* A736-derived ILA regulates ChAT-expressing neurons and immune responses of DSS-induced mice

We next identified specific tryptophan-derived metabolites responsible for these effects. Metabolomic profiling of feces and *L. plantarum* A736 supernatant revealed a significant elevation in indole-3-lactic acid (ILA) - a key tryptophan metabolite (**Figure 5A-5B**). To validate these findings, we quantified tryptophan-related metabolites via liquid chromatography-MS (LC-MS). *L. plantarum* A736 administration markedly elevated fecal ILA levels, an effect abolished under a low-tryptophan diet (**Figure 5C-5D**). In contrast, the remaining tryptophan metabolites had no obvious changes following *L. plantarum* A736 supplementation **(Figure S7G-S7H)**. The ILA production and a corresponding decrease in tryptophan levels were also observed in the *L. plantarum* A736 supernatant following culture in MRS medium (**Figure 5E and Figure S7I)**. These findings demonstrate that *L. plantarum* A736 could metabolize tryptophan into ILA. Whole-genome sequencing of L. plantarum A736 identified ldhA, which encodes 2-hydroxyacid dehydrogenase—a key enzyme in ILA biosynthesis **(Figure S7J)**. We further confirmed the production of ILA by *L. plantarum* A736 and in peptone-tryptone-tryptophan (PTT) media accompanied by significantly reduced tryptophan levels, suggesting the ability of *L. plantarum* A736 for converting tryptophan to ILA (**Figure 5F and Figure S7K)**. Similarly, UC patients showed significantly lower fecal ILA levels compared to healthy controls. Pearson correlation analysis revealed negative associations between fecal ILA concentrations and clinical disease indicators **(Figure S7L-7M)**. Together, these results suggest that *L. plantarum* A736-derived ILA may contribute to colitis amelioration.

**Figure 5.**
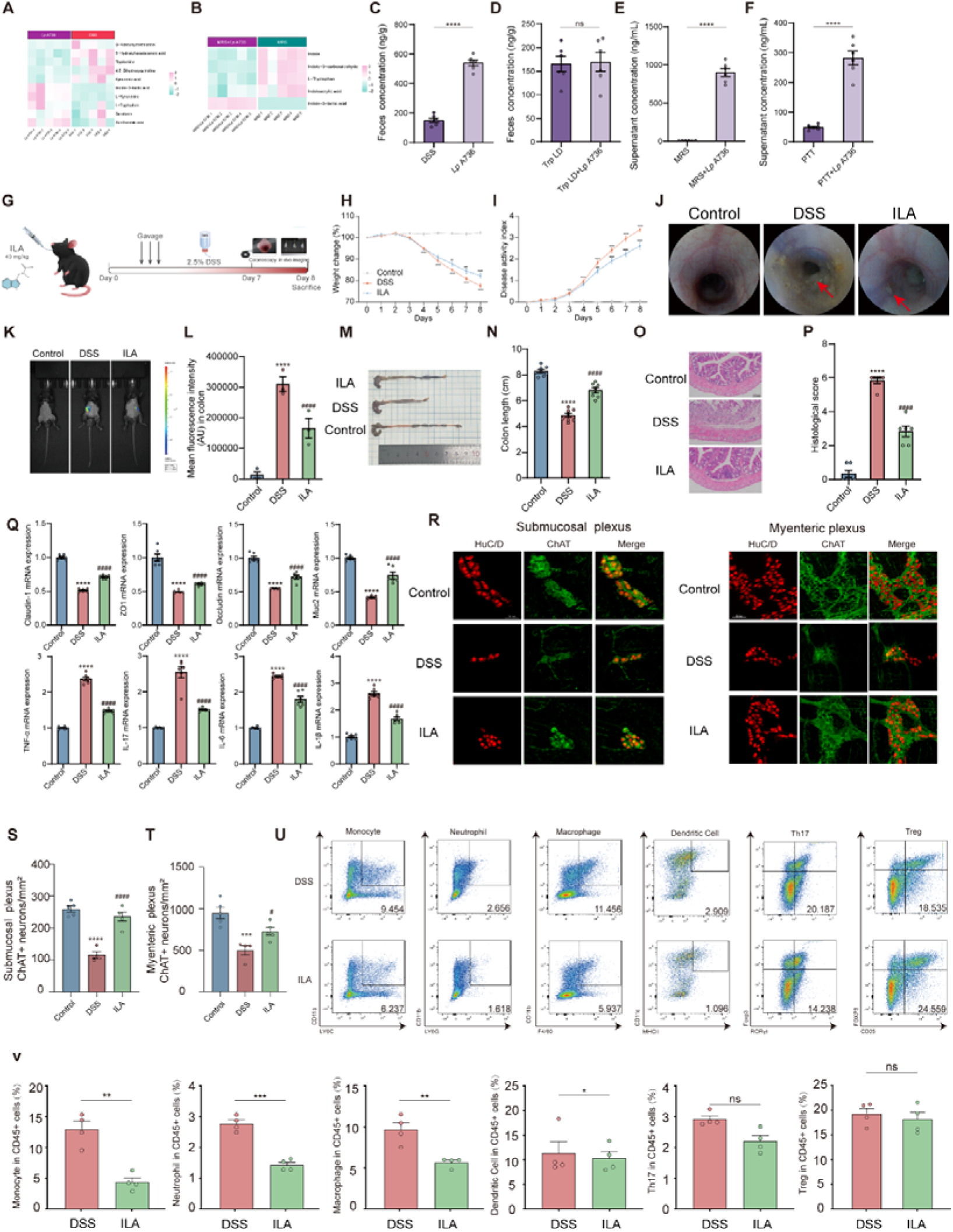
*L. plantarum* A736-derived ILA regulates ChAT-expressing neurons and and immune responses in DSS-induced mice. (A) Heatmap displaying the enrichment of tryptophan-related fecal metabolites in mice between DSS and *Lp* A736 (n=5). (B) Heatmap displaying the enrichment of tryptophan-related metabolites between MRS and MRS *+Lp* A736 (n=5). (C) Concentrations of indole-3-lactic acid (ILA) in feces from mice between DSS and *Lp* A736 (n=6). (D) Concentrations of indole-3-lactic acid (ILA) in feces from mice treated with *L. plantarum* A736 or PBS during low-tryptophan diet between DSS and *Lp* A736 (n=6). (E) ILA production by *L. plantarum* A736 in MRS medium (n=6). (F) ILA production by *L. plantarum* A736 in peptone-tryptone-tryptophan (PTT) medium (n=6). (G) Schematic diagram for ILA treatment. (H) Body weight change (n=8). (I) Disease activity index (DAI) score (n=8). (J) Representative colonoscopy images (n=3). (K-L) Representative image captured by an in vivo imaging system and quantification (n=3). (M) Representative photographs of colon. (N) Colon length (n=8). (O and P) Representative image of hematoxylin and eosin (H&E)-staining sections in colon and histological scores (n=6). Scale bar, 50 μm. (Q) The mRNA expression of gut barrier-related proteins (Claudin-1, ZO1, Occludin and Muc2) and pro-inflammatory cytokines (TNF-α, IL-17, IL-6 and IL-1β) in colonic tissues evaluted by quantitative real-time PCR (n=6). (R) ChAT (green) and HuC/D (red) immunostaining of colonic submucosal and myenteric plexus (n=4-5). Scale bar, 36.8 μm. (S-T) Quantification of ChAT^+^ neurons in both submucosal (S) and myenteric plexus (T) of the colon (n=4-5). (U) Representative flow cytometry plots for monocytes, macrophages, neutrophils, dendritic cells, Th17 and Treg cells from colonic lamina propria (cLP). (V) The percentage of monocytes, macrophages, neutrophils, dendritic cells, Th17 and Treg cells (n=4). Control group (Control), DSS group (DSS), *L. plantarum* A736 treatment group (*Lp* A736) and ILA treatment group (ILA). Low-tryptophan (Trp) diet (Trp LD), low-tryptophan (Trp) diet and *L. plantarum* A736 (Trp LD+*Lp* A736). Original culture medium (Man-Rogosa-Sharpe, MRS) and *L. plantarum* A736 culture supernatant (MRS+*Lp* A736). Peptone-tryptone-tryptophan medium (PTT) and *L. plantarum* A736 supernatants in peptone-tryptone-tryptophan medium (PTT+ *Lp* A736). Mean ± SEM shown. Data are determined by two-way ANOVA with Tukey’s multiple comparison test (H-I), one-way ANOVA with Tukey’s multiple comparison test (M-T) or unpaired Student’s t-test (C-F and V). ns, not significant, *p < 0.05, **p < 0.01, ***p < 0.001, and ****p < 0.0001, compared with the Control group, ^#^p<0.05, ^##^p<0.01, ^###^p < 0.001, and ^####^p < 0.0001, compared with the DSS group. See also Figures S7.

To directly determine the impact of ILA on colitis, we treated mice with or without ILA during DSS exposure and assessed changes in weight loss, disease activity index, histological score, colon length, endoscopy findings, and in vivo imaging. ILA treatment had effects similar to those of *L. plantarum* A736, ameliorating the disease phenotypes as described above (**Figure 5G-5P**). Furthermore, ILA reduced intestinal permeability and downregulated the expression of colonic pro-inflammatory mediators (**Figure 5Q**). Notably, oral ILA administration restored the number of detectable ChAT-expressing neurons in whole-mount-stained submucosal and myenteric plexuses, which were diminished in DSS-induced colitis (**Figure 5R-5T**). In the colonic lamina propria, ILA treatment during DSS colitis significantly reduced the proportions of monocytes, macrophages, neutrophils, and Th17 cells, but not dendritic cells or Treg cells. These findings suggest that L. plantarum A736-derived ILA acts as a microbial signal contributing to colitis remission, potentially through modulation of enteric ChAT-expressing neurons and intestinal immunity (**Figure 5U-5V**).

##### *L. plantarum* A736-sensing is dependent on ChAT-expressing neurons to intestinal immunity

To investigate the functional role of cholinergic neurons in *L. plantarum* A736-mediated immune regulation, we performed targeted ablation of ChAT-expressing neurons in ChAT-Cre mice through colonic administration of a Cre-dependent diphtheria toxin A (DTA) virus (rAAV-EF1α-DIO-DTA-WPRE-hGH polyA; ChAT^AAV-DTA^) (**Figure 6A-6B**). Control mice received the corresponding EGFP-expressing virus (rAAV-EF1α-DIO-EGFP-WPRE-hGH polyA; ChAT ^AAV-EGFP^). Whole-mount immunofluorescence analysis of isolated myenteric plexuses revealed a significant reduction in ChAT^+^ neurons in ChAT ^AAV-DTA^ mice compared to ChAT ^AAV-EGFP^ controls, confirming efficient neuronal ablation **(Figure S7N)**.

**Figure 6.**
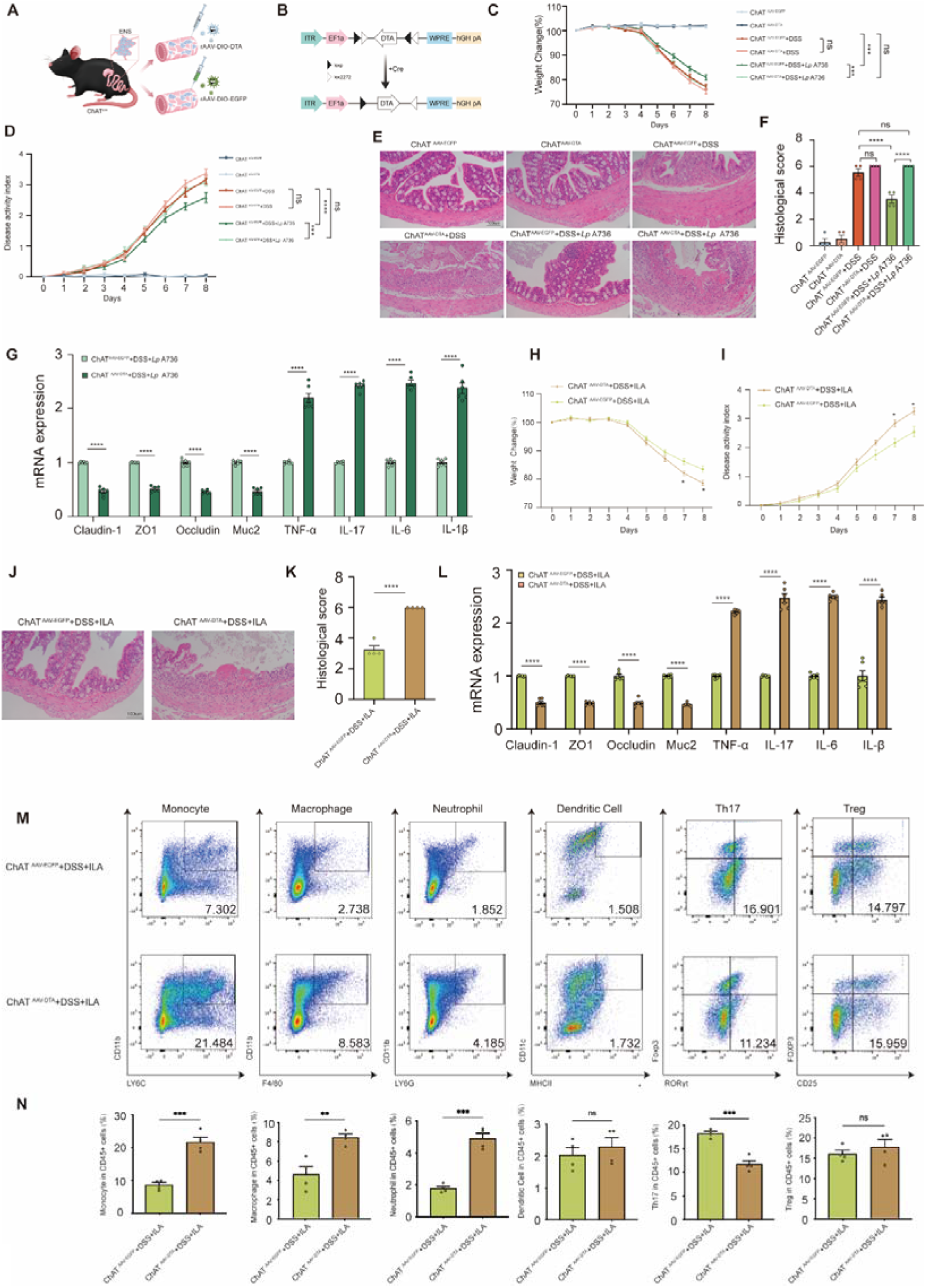
ChAT-expressing neurons ablation mediate *L. plantarum* A736-regulated intestinal immunity in colitis. (A) Schematic diagram illustrating the intracolonic injections of AAV-encoding EGFP or DTA into the ChAT-iCre mice. (B) Illustration of the rAAV-EF1α-DIO-DTA-WPRE-hGH polyA construct expressing diphtheria toxin subunit A (DTA) in a Cre-dependent configuration to cause selective cell death in Cre-expressing neurons. (C-G) Body weight change (n=8) (C), Disease activity index (DAI) score (D), representative image of hematoxylin and eosin (H&E)-staining sections in colon and histological scores (n=4). Scale bar, 50 μm (E and F), the mRNA expression of gut barrier-related proteins (Claudin-1, ZO1, Occludin and Muc2) and pro-inflammatory cytokines (TNF-α, IL-17, IL-6 and IL-1β) in colonic tissues evaluted by quantitative real-time PCR (n=6) (G) from ChAT ^AAV-EGFP^, ChAT^AAV-DTA^, ChAT ^AAV-EGFP^+DSS, ChAT^AAV-DTA^+ DSS, ChAT ^AAV-EGFP^+ DSS+*Lp* A736 and ChAT^AAV-DTA^+ DSS+*Lp* A736 mice. (H-N) Body weight change (n=8) (H), Disease activity index (DAI) score (n=8) (I), representative image of hematoxylin and eosin (H&E)-staining sections in colon and histological scores (n=4). Scale bar, 50 μm (J and K), the mRNA expression of gut barrier-related proteins (Claudin-1, ZO1, Occludin and Muc2) and pro-inflammatory cytokines (TNF-α, IL-17, IL-6 and IL-1β) in colonic tissues evaluted by quantitative real-time PCR (n=6) (L), Representative flow cytometry plots for monocytes, macrophages, neutrophils, dendritic cells, Th17 and Treg cells from colonic lamina propria (cLP) (M), the percentage of monocytes, macrophages, neutrophils, dendritic cells, Th17 and Treg cells (n=4) (N) from ChAT ^AAV-EGFP^+DSS+ILA and ChAT^AAV-DTA^+DSS+ILA mice. PBS-treated ChAT-iCre mice infected with AAV-encoding EGFP viruses (ChAT ^AAV-EGFP^), PBS-treated ChAT-iCre mice infected with AAV-encoding DTA viruses (ChAT^AAV-DTA^), DSS-treated ChAT-iCre mice infected with AAV-encoding EGFP viruses (ChAT^AAV-EGFP^+DSS), DSS-treated ChAT-iCre mice infected with AAV-encoding DTA viruses (ChAT^AAV-DTA^+ DSS), DSS-treated ChAT-iCre mice infected with AAV-encoding EGFP viruses and orally administered *L. plantarum* A736 (ChAT ^AAV-EGFP^+ DSS+*Lp* A736), DSS-treated ChAT-iCre mice infected with AAV-encoding DTA viruses and orally administered *L. plantarum* A736 (ChAT^AAV-DTA^+ DSS+*Lp* A736), DSS-treated ChAT-iCre mice infected with AAV-encoding EGFP viruses and orally administered ILA (ChAT^AAV-EGFP^+ DSS+ ILA), DSS-treated ChAT-iCre mice infected with AAV-encoding DTA viruses and orally administered ILA (ChAT^AAV-DTA^+ DSS+ ILA). Data are determined by one-way ANOVA with Tukey’s multiple comparison test (C-G) or unpaired Student’s t-test (H-N). ns, not significant, *p < 0.05, **p < 0.01, ***p < 0.001, and ****p < 0.0001. See also Figures S7.

Critically, ablation of ChAT-expressing neurons completely abolished the protective effects of *L. plantarum* A736 against DSS-induced colitis, as evidenced by significantly attenuated improvements in weight loss, disease activity index, histological scores, colon shortening, and expression of mucosal barrier-related proteins and pro-inflammatory mediators in colonic tissues (**Figure 6C-G**). Similarly, neuronal ablation eliminated the beneficial effects of *L. plantarum* A736-derived ILA (**Figure 6H-L**). Flow cytometry revealed distinct immune cell infiltration patterns in the colonic lamina propria (cLP) of ILA-treated ChAT ^AAV-DTA^ mice versus ChAT ^AAV-EGFP^ controls during DSS colitis (**Figure 6M-N**), demonstrating that ChAT^+^ neurons are neurons are necessary and sufficient for L. plantarum A736/ILA-mediated immunoregulation. These findings establish ChAT-expressing neurons as critical regulatory nodes that mediate the intestinal immune-modulatory effects of both *L. plantarum* A736 and its derived metabolite ILA.

#### Discussion

Previously, UC patients and DSS-induced colitis mice exhibit significantly lower *Lactobacillus* genus abundance.^66,72,73^ Beneficial effect to clinical indicators and inflammation suppression was associated with increased *Lactobacillus*.^66,73^ Here, we further validated that the abundance of *Lactobacillus* was negatively correlated with the disease phenotypes of UC patients and colitis mice, which corroborates previous studies. This motivated us to explore *Lactobacillus* strains as a potential candidate for microbial-based therapeutic strategy to UC. However, the efficacy of probiotics is strain-dependent. Hence, based on screening *Lactobacillus* isolates from fermented foods both in vitro and in vivo experiments, we found that *L. plantarum* A736 effectively rescued colitis-related phenotypes. In addition, 5-aminosalicylic acid (5-ASA), a therapy for UC remission in clinical guideline,^68^ was also used to confirm the efficacy of *L. plantarum* A736. We observed comparable effects in inhibiting inflammation and restoring gut barrier damage, whereas *L. plantarum* A736 exhibited more superiorities, especially in alleviating weight loss and increased disease activity index. These outcomes provide compelling evidence for *L. plantarum* A736 as a potential alternative therapy in IBD treatment. Extensive clinical research involving diverse ethnic populations among the general global public is required to validate the protective role for *L. plantarum* A736 in the progression of IBD.

ILA, a tryptophan-indole metabolite derived from bacteria, can act as a ligand to activate aryl hydrocarbon receptor (AhR), which has been implicated in gastrointestinal disorders.^74–76^ Recent clinical studies suggest reduced ILA levels in both IBD patients^77^ and colitis mice^73^ with additional evidence demonstrating that oral administration of ILA ameliorates colitis progression in mice model. For example, ILA relieves colitis indicators by inhibiting epithelial CCL2/7 production to decrease the accumulation of inflammatory macrophages.^51^ Likewise, *L. plantarum*-derived ILA markedly alleviates pathological symptoms in acute-colitis mice.^78^ Indeed, ILA produced by *L. johnsonii* contributes to the mitigation of colitis through AhR signaling pathway.^73^ In both DSS-induced and IL-10^−/−^ spontaneous colitis models, ILA is capable to inhibit gut inflammation and alter the composition of microbiota, thus enhancing the abundance of tryptophan-metabolizing bacteria for the production of other indole derivatives.^79^ These findings, in concert with our work, support that *L. plantarum* A736 ameliorate colitis progression mainly by its metabolite ILA. Generally, a variety of *Lactobacillus* species including *Lactobacillus reuteri*, *Lactobacillus johnsonii*, and *Lactobacillus plantarum* possesses the ability convert tryptophan into ILA,^75,80^ which reveal several key enzymes for the ILA biosynthesis, such as aromatic amino acid aminotransferase (ArAT)^45,79,81^ and D-2-hydroxyacid dehydrogenase (ldhA).^42,82^ In line with but extending further, we noted the existence of LdhA in the genome of *L. plantarum* A736 for the very first time and further confirmed the ability of *L. plantarum* A736 for metabolizing tryptophan to produce ILA. Specifically, our findings indicated that L. *L. plantarum* A736-released metabolite ILA was dependent on dietary tryptophan, because ILA production were significantly reduced in low tryptophan diet. However, the role of ldhA enzyme involved in ILA production by *L. plantarum* A736 still require further confirmation using a mutant strain that lacks the 2-hydroxyacid dehydrogenase(*L. plantarum* A736 △ldhA).

Abnormalities of ENS structure and function have been described in IBD. Clinical observations have documented a lower number of enteric neurons in IBD patients,^13^ especially catecholaminergic and cholinergic neurons.^14–15^ Of note, cholinergic neurons^14^ release the neurotransmitter ACh to communicate with other cells that is attracting considerable interest as a major subset of enteric neurons expressing ChAT.^12,16^ Another animal studies with IBD also note diminished the expression of ChAT and ACh levels.^14,19–21^ As previously mentioned, ACh is considered as the most important neurotransmitter in the ENS, which are involved in regulating inflammatory response through ACh receptors expressing on various immune cells, including mAChRs and nAChRs respectively. ACh has been confirmed to protect against colitis by nAChR/ERK pathway to promote IL-10 production in monocytic myeloid-derived suppressor cells (M-MDSCs).^14^ Also, nicotine treatment rescued colitis severity after DSS exposure, due to restoring nicotinic acetylcholine receptor signaling levels. In both experimental colitis models induced by 2,4-dinitrobenzensulfonic acid (DNBS) or DSS, correction of ENS defects in TLR2^-/-^ mice by glial cell line-derived neurotrophic factor (GDNF) administration massively ameliorates body weight change and reduce inflammatory response.^83,84^ Recently, optogenetic activation of enteric cholinergic neurons decrease proinflammatory cytokine expression and relieve colitis severity. Similarly, activation of cholinergic neurons with blue light stimulation exhibits a decrease of IL-1β expression from muscularis macrophages in vitro.^17^ For the very first time, our data revealed that *L. plantarum* A736 and its metabolite ILA is able to balance immune responses by cholinergic neurons, thereby ameliorating colitis. Our work also added to the evidence of neuroimmune regulation. Overall, we highlighted the integral role of cholinergic neurons in neuroimmune regulation, supporting that *L. plantarum* A736 treatment to restore enteric cholinergic may represent a novel target to probiotic-based therapies in gastrointestinal disorders. Currently, direct evidence regarding the involvement of probiotics and indole derivatives in the regulation of ENS is currently very scarce and remains in the stage of data accumulation, while our study have focused on these aspects. Moreover, hardly any studies have suggested that ILA directly impacts cholinergic neurons, and the detailed molecular mechanism remains uncertain, although ILA have been identified to possess AhR-activating properties.^74^ But our work only confirmed that *L. plantarum* A736-derived ILA impacts cholinergic neurons. We speculate *L. plantarum* A736-released ILA might contribute to activate AHR signaling in enteric cholinergic neurons and further influence AhR-dependent transcriptional program, based on a previous study suggesting that AhR is highly expressed in colonic HuC/D^+^ enteric neurons including ChAT-expressing neurons,^31^ which indicates that these neurons represent the main target of AhR ligands in this gut layer.

In conclusion, we demonstrate a previously unrevealed role of ILA-producing *L. plantarum* A736 on regulating unbalanced intestinal immunity via enteric ChAT-expressing neurons, resulting in the alleviation of colitis symptoms. These findings contribute to improving our understanding of the neuroimmune mechanism underlying IBD pathogenesis and bacterial-mediated immunoregulation. Furthermore, we provide a rationale for possible future clinical applications of *L. plantarum A736* and its metabolite ILA in microbial-based therapeutic strategies for gastrointestinal disorders, including but not limited to IBD.

#### Limitations of the study

There are some limitations regarding to our study. Firstly, although we show the existence of ldhA in the genome of *L. plantarum* A736, the crucial role of ldhA to encode enzyme for ILA production in this strain need be better confirmed by generating a genetically modified *L. plantarum* A736 (*L. plantarum* A736 △ldhA) that lacks 2-hydroxyacid dehydrogenase, which enable us to mechanistically dissect the specific role of *L. plantarum* A736-synthesized ILA for relieving colitis. Furthermore, we observed that ILA, a molecule with AhR ligand activity, regulated intestinal immunity functions via cholinergic neurons. However, the exact molecular mechanism by which ILA affects cholinergic neurons remain incompletely addressed. Whether ILA activates AhR signaling and acts downstream of neuronal AhR within cholinergic neurons should be clarified by generating new conditional knockout mice with cholinergic-neuron-specific deletion of AhR in future studies. Additionally, we demonstrated that acetylcholine produced by cholinergic neurons was promoted by *L. plantarum* A736, and previous studies indicated that ACh exerted immunoregulatory capacity through ACh receptors on immune cells. But detailed downstream signaling pathways involved in neuroimmune regulatory mechanism by *L. plantarum* A736 treatment remains to be characterized. Lastly, a randomized placebo-controlled clinical trial is needed to explore the efficacy of *L. plantarum* A736 for the treatment of IBD.

## Acknowledgements

This work was financially supported by the National Natural Science Foundation of China (No. 32472354), National Natural Science Foundation of China (No. 32101968) and Shaanxi Provincial Natural Science Basic Research Program (No. 2024JC-YBQN-0094). We thank the Life Science Research Core Services, NWAFU (Zhou Min, Yang Chenghui, Liu Xiaorui, Liu Yao) for technical support.

---

**Figure S1.**
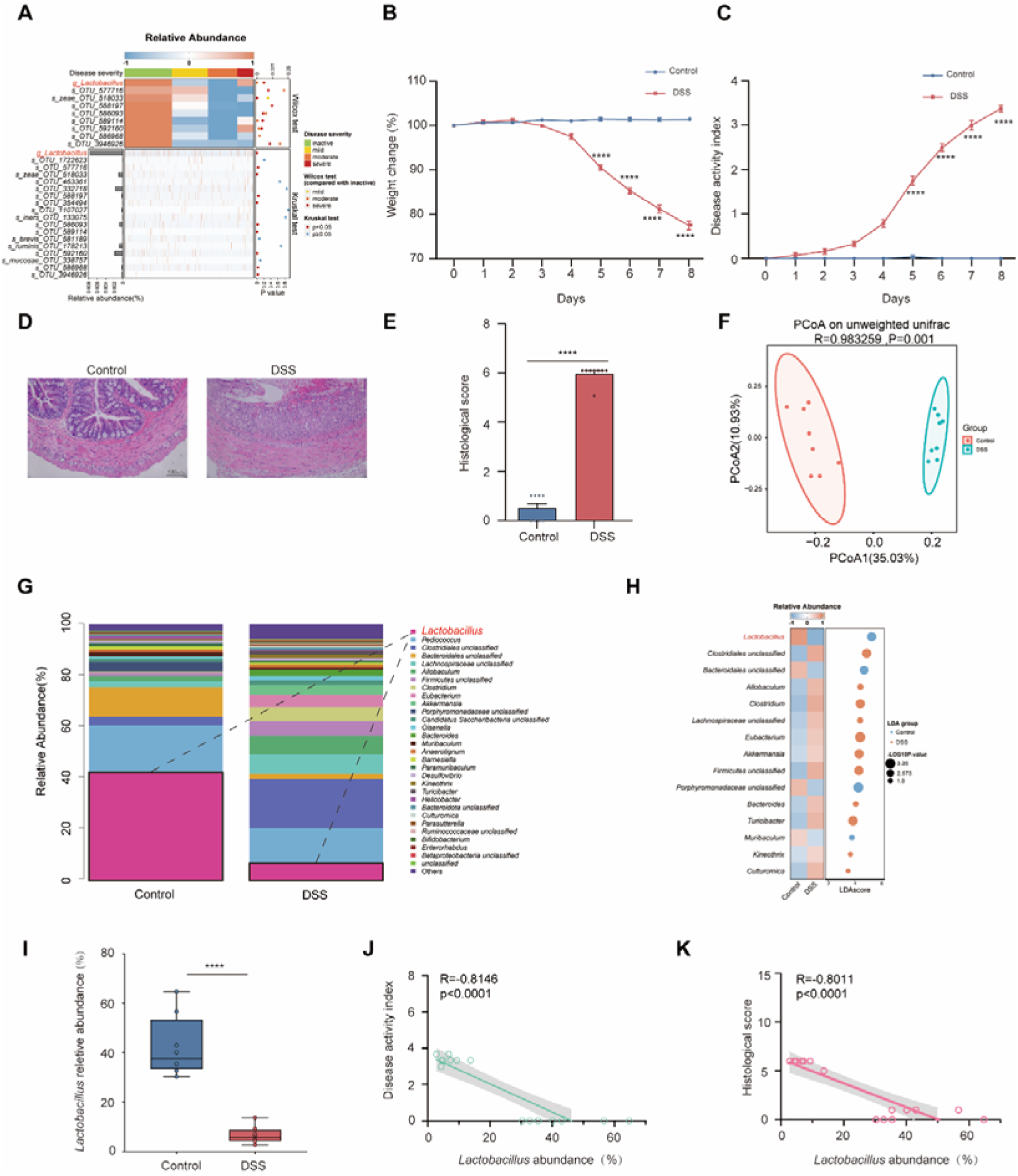
The abundance of *Lactobacillus* is reduced in UC patients and DSS-induced mice, related to Figure 1. (A) The abundance of *Lactobacillus* genus from UC patients in diverse disease course by a previous report.^67^ (B) Body weight change. (C) Disease activity index (DAI). (D) Representative image of hematoxylin and eosin (H&E)-staining sections in colon. Scale bar, 50 μm. (E) Histological scores. (F) Principal co-ordinates analysis (PCoA) of stool samples based on unweighted unifrac distances by 16S rRNA sequencing. ANOSIM (analysis of similarities), R = 0.983259, p = 0.001. (G) Averaged relative abundance of bacteria at the genus level from Control and DSS. (H) Linear discriminative analysis (LDA) score of differentially enriched bacterial genera measured by Linear discriminant analysis Effect Size (LEfSe) analysis between Control and DSS. (I) Relative abundance of *Lactobacillus* genus from Control and DSS in mice. (J-K) Correlation analysis of *Lactobacillus* abundance with DAI (J) and histological scores (K). Control group (Control, n=8) DSS group (DSS, n=8). Mean ± SEM shown. ns, not significant, *p < 0.05, **p < 0.01, ***p < 0.001, and ****p < 0.0001, determined by unpaired Student’s t-test.

**Figure S2.**
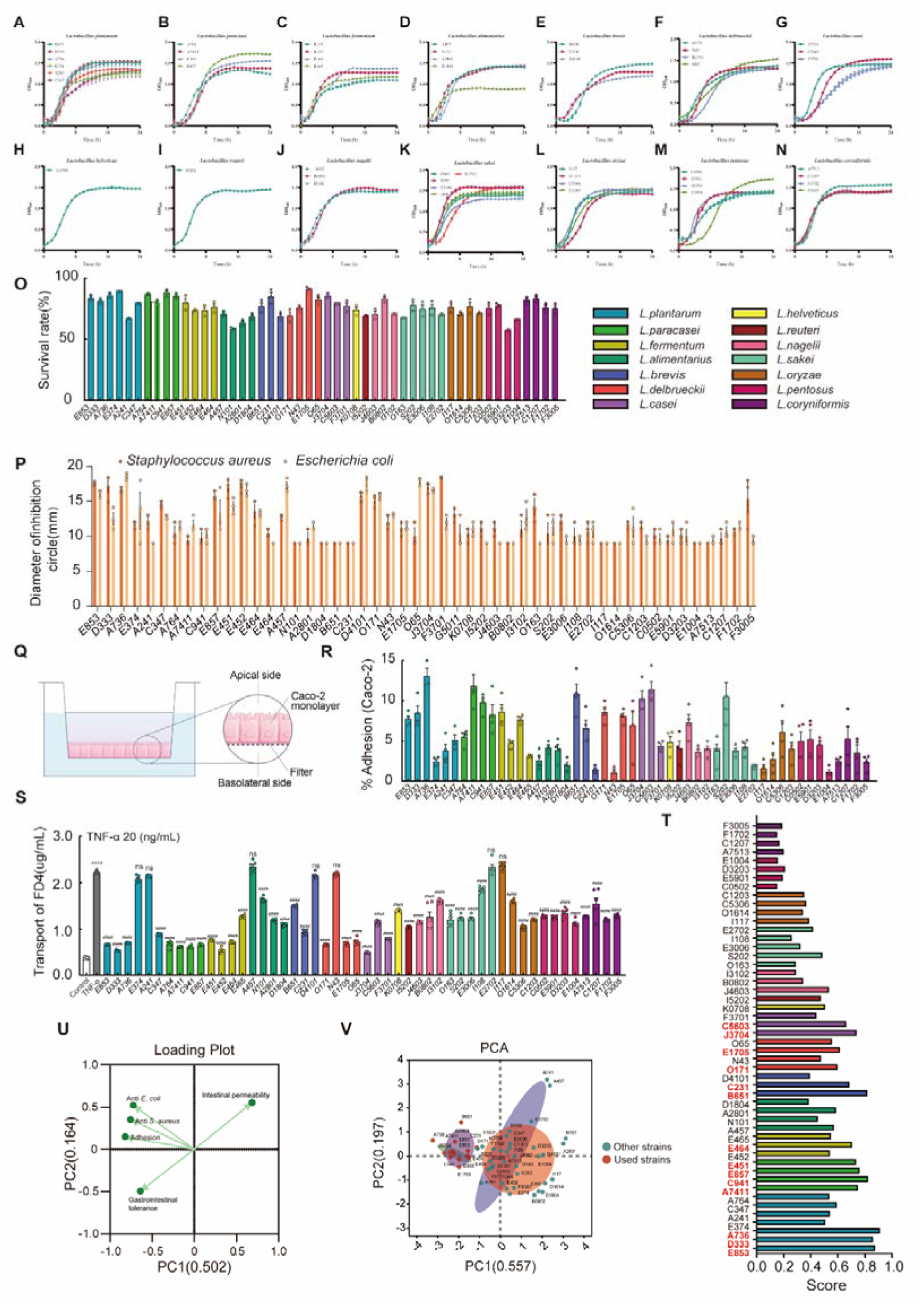
Physiological characteristics of *Lactobacillus* strains in vitro, related to Figure 2. (A-N) Growth curve of *L. plantarum* (A), *L. paracasei* (B), *L.fermentum* (C), *L. alimentarius* (D), *L. brevis* (E), *L. delbrueckii* (F), *L.casei* (G), *L.helveticus* (H), *L.reuteri* (I), *L. nagelii* (J), *L. sakei* (K), *L.oryzae* (L), *L.pentosus* (M), and *L.coryn iformis* (N) (n=3). (O) The survival rate of *Lactobacillus* strains following simulated gastrointestinal fluids. (P) Antibacterial activity. (Q) Diagram of a Caco-2 monolayer. (R) Adhesion of the *Lactobacillus* strains with Caco-2 cell monolayers (MOE 10, n=3). (S) 4 kDa FITC-dextran paracellular transport from the apical to basolateral compartment on a Caco-2 cell monolayer pretreated with *Lactobacillus* strains (MOE; 10, 24 h) before TNFα exposure (20 ng/mL, 24 h, n=3). (T) Loading plot of principal component analysis (PCA). (U) Score plot of PCA based on the survival rate, antibacterial activity, adhesion and intestinal permeability of *Lactobacillus* strains in vitro. (V) Fuzzy synthetic evaluation (FSE) analysis. Mean ± SEM shown. Data are determined by one-way ANOVA with Tukey’s multiple comparison test. ns, not significant, *p < 0.05, **p < 0.01, ***p < 0.001, and ****p < 0.0001, compared with the Control group, ^#^p<0.05, ^##^p<0.01, ^###^p < 0.001, and ^####^p < 0.0001, compared with the TNFα group.

**Figure S3.**
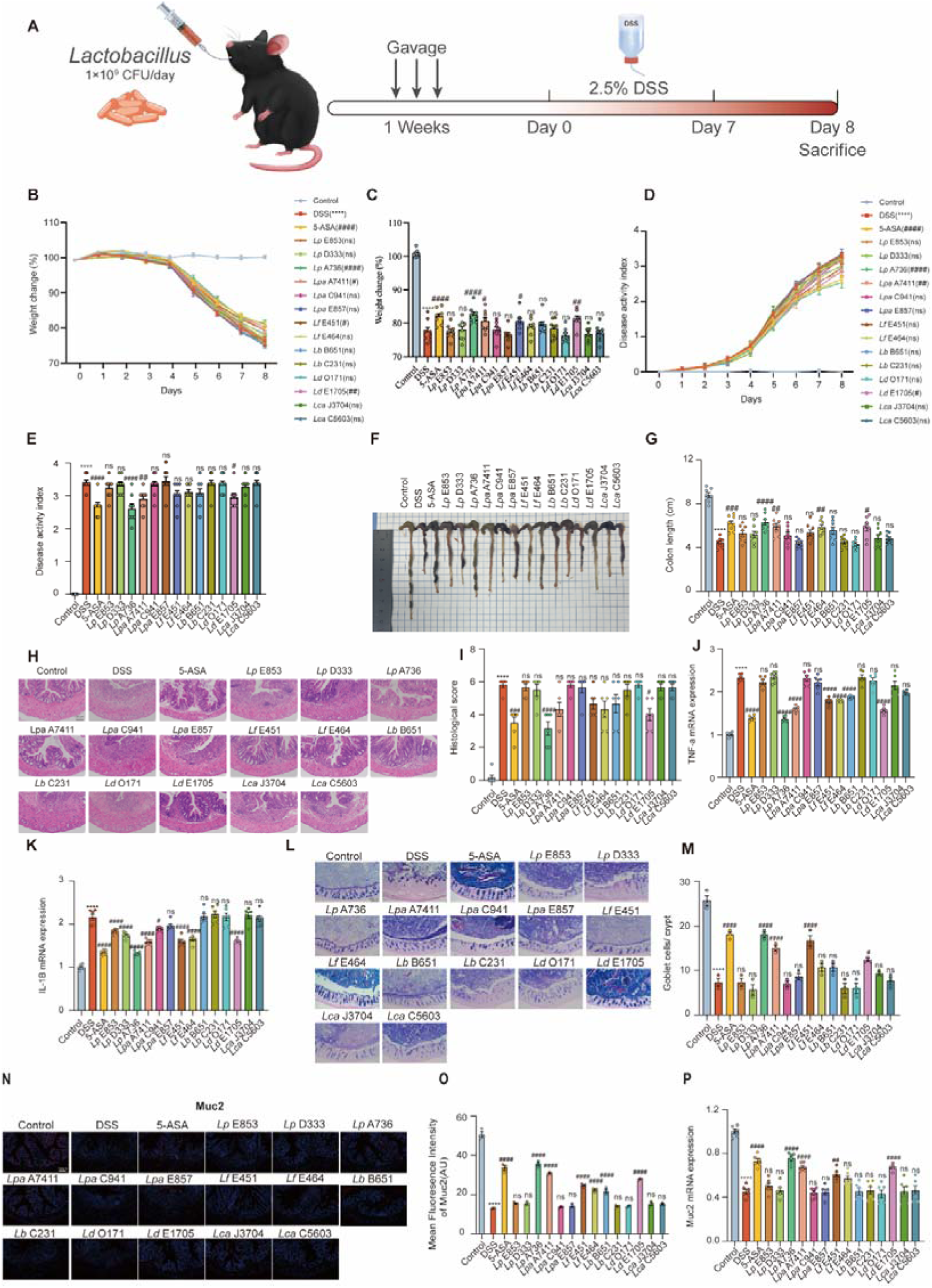
*Lactobacillus* strains ameliorates colitis symptoms, inflammation and mucus barrier damage, related to Figure 2. (A) Experimental diagram for *Lactobacillus* strains administration schedule. (B) Body weight change (n=8). (C) Body weight change at day 8 after DSS induction (n=8). (D) Disease activity index (DAI) score (n=8). (E) DAI at day 8 after DSS induction (n=8). (F) Representative photographs of colon. (G) Colon length (n=8). (H and I) Representative image of hematoxylin and eosin (H&E)-staining sections in colon and histological scores (n=6). Scale bar, 50 μm. (J-K) Relative gene expression of proinflammatory cytokines including TNF-α (J) and IL-1β (K) in colonic tissues accessed by quantitative real-time PCR. (L-M) AB/PAS staining of the distal colon and elevation of goblet cells per crypt (n=3). Scale bar, 50 μm. (N-O) Representative immunostaining images and quantification of Muc2 in the colon (n=3). Scale bar, 50 μm. (P) Relative gene expression of Muc2 in colonic tissues accessed by quantitative real-time PCR. Data are determined by two-way ANOVA with Tukey’s multiple comparison test (B-E) or one-way ANOVA with Tukey’s multiple comparison test (G-O). ns, not significant, *p < 0.05, **p < 0.01, ***p < 0.001, and ****p < 0.0001, compared with the Control group, ^#^p<0.05, ^##^p<0.01, ^###^p < 0.001, and ^####^p < 0.0001, compared with the DSS group.

**Figure S4.**
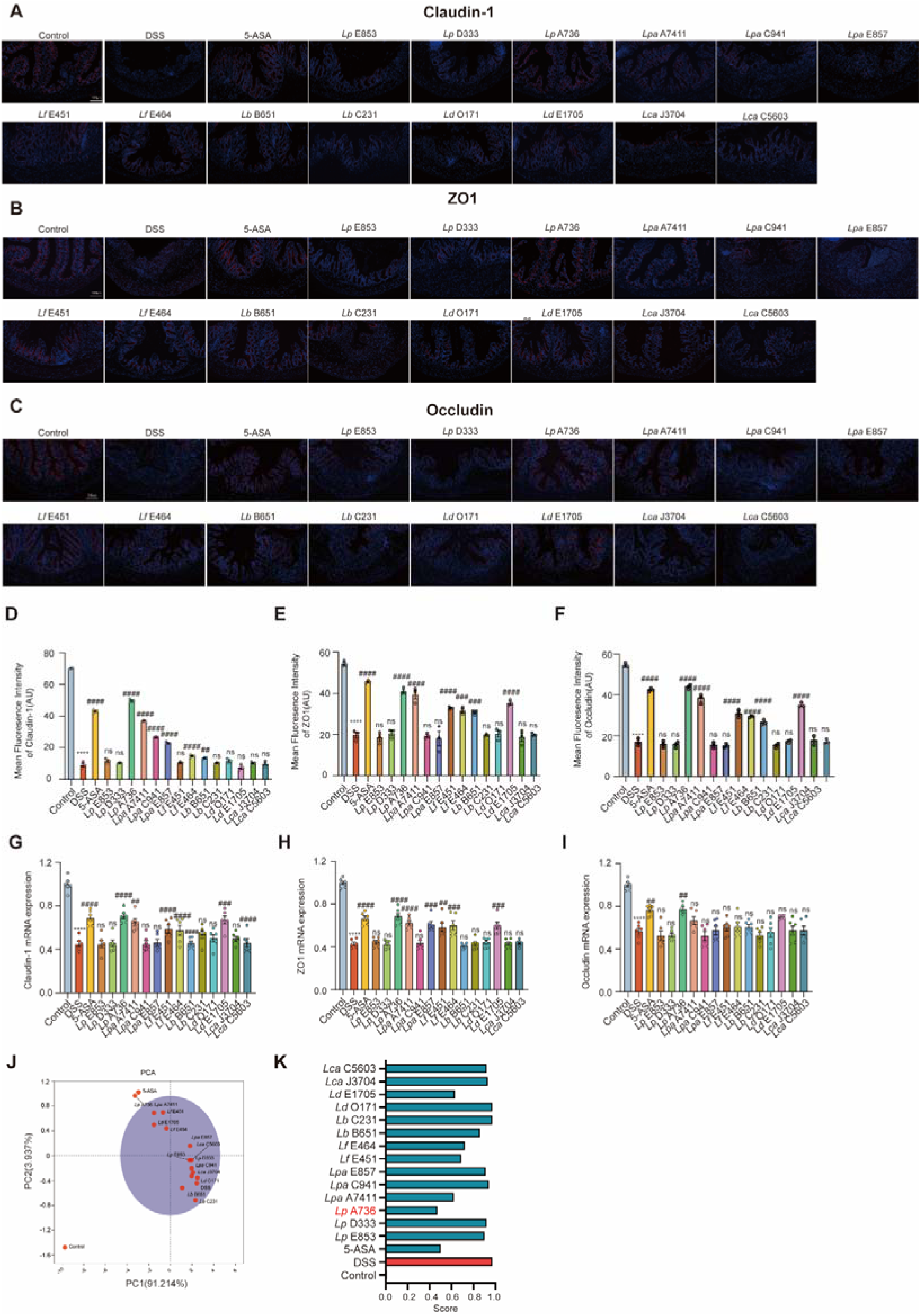
*Lactobacillus* strains ameliorates resulting epithelial barrier damage, related to Figure 2. (A-F) Representative immunofluorescence staining images and quantification of Claudin-1, ZO1 and Occludin in the colon (n=3). Scale bar, 50 μm. (G-I) Relative gene expression of Claudin-1(G), ZO1(H) and Occludin (I) in colonic tissues accessed by quantitative real-time PCR. (J) Principal component analysis (PCA) based on the body weight change, disease activity index, colon length, histological scores, goblet cells, the mRNA expression of TNF-α, IL-1β, Muc2, Claudin-1, ZO1 and Occludin in colon tissues. (K) Fuzzy synthetic evaluation (FSE) analysis. Data are determined by one-way ANOVA with Tukey’s multiple comparison test. ns, not significant, *p < 0.05, **p < 0.01, ***p < 0.001, and ****p < 0.0001, compared with the Control group, ^#^p<0.05, ^##^p<0.01, ^###^p < 0.001, and ^####^p < 0.0001, compared with the DSS group. See also Figures S2, S3 and S4, Table S1.

**Figure S5.**
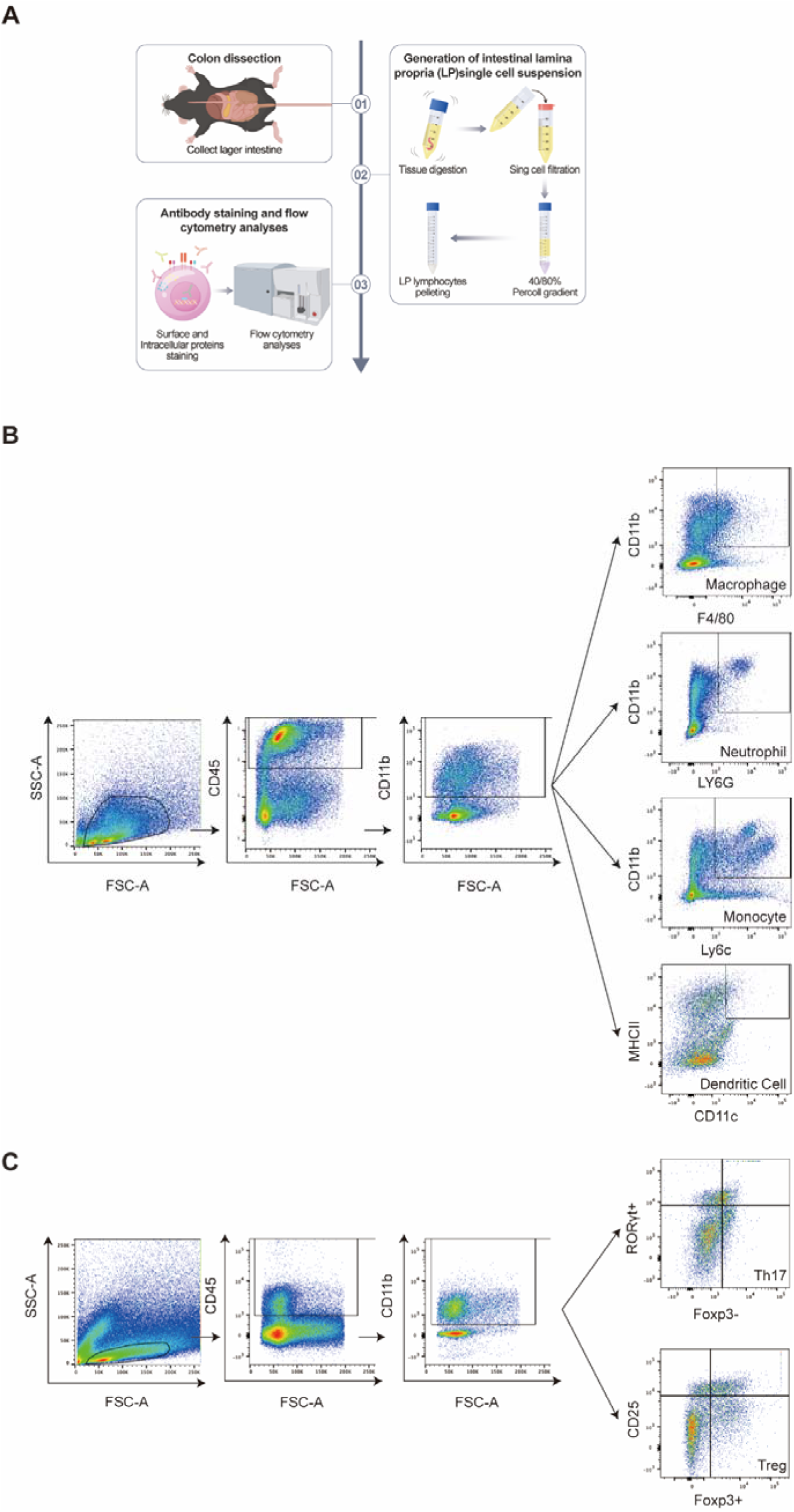
*L. plantarum* A736 regulates intestinal immune, related to Figure 2. (A) Experimental diagram for isolation of immune cells from colonic lamina propria (cLP). (B) The gating strategies for flow cytometry analysis of monocytes, macrophages, neutrophils and dendritic cells from cLP. (C) The gating strategies for flow cytometry analysis of Th17 and Treg cells from cLP.

**Figure S6.**
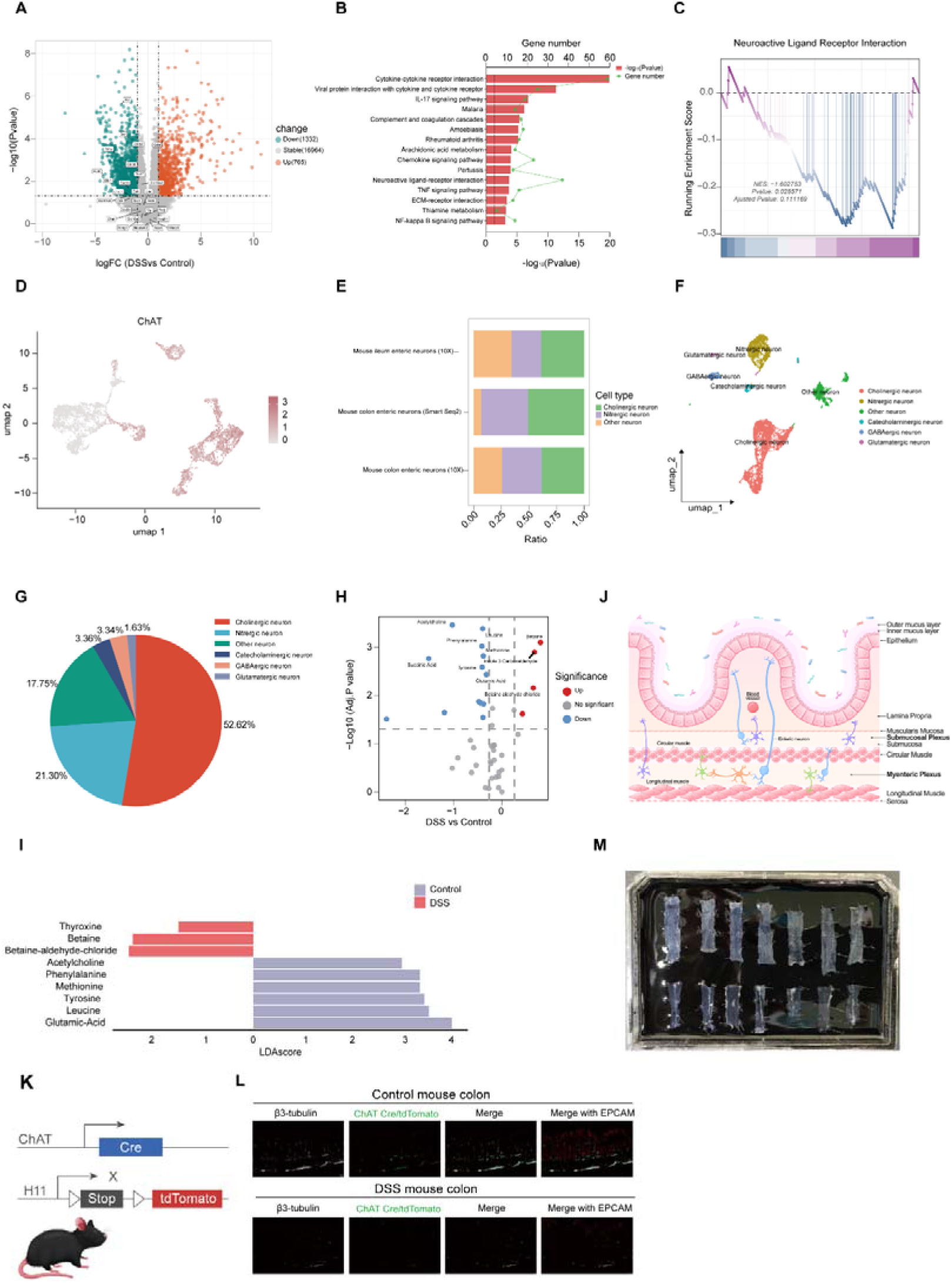
*L. plantarum* A736 impacts neuroactive signaling pathways, neurotransmitter production and ChAT-expressing neurons, related to Figure 3. (A) Volcano plot displaying differentially expressed genes in colon tissues between DSS and Control (n=8). (B) KEGG analysis demonstrating distinct differences in pathways between DSS and Control (n=8). (C) GSEA(n=8). (D-E) The distribution of ChAT (D) and the proportion of major enteric neurons classes in colon or ileum from mice (E) by a published study.^12^ (F-G) UMAP (F) and pie chart (G) displaying the subclustering of enteric neurons or the percentage of enteric neurons classes respectively by a published study.^71^ (H) Volcano plot showing differential neurotransmitters in colon tissues between DSS and Control (n=8). (I) Linear discriminative analysis (LDA) score of differentially enriched neurotransmitters from LEfSe analysis between DSS and Control (n=8). (J) Schematic of the ENS in the colon. (K) Schematic showing ChAT-Cre/tdTomato reporter mice. (L) Immunostaining of colonic sections from ChAT-Cre/tdTomato reporter mice stained for pan-neuronal marker βIII-tubulin (green) and epithelia marker EpCam1 (white) in Control and DSS group. Scale bar, 50 μm. (M) Sylgard plate with multiple pinned fragments of colonic myenteric and submucosal plexus. Control group (Control), DSS group (DSS) and *L. plantarum* A736 treatment group (*Lp* A736).

**Figure S7.**
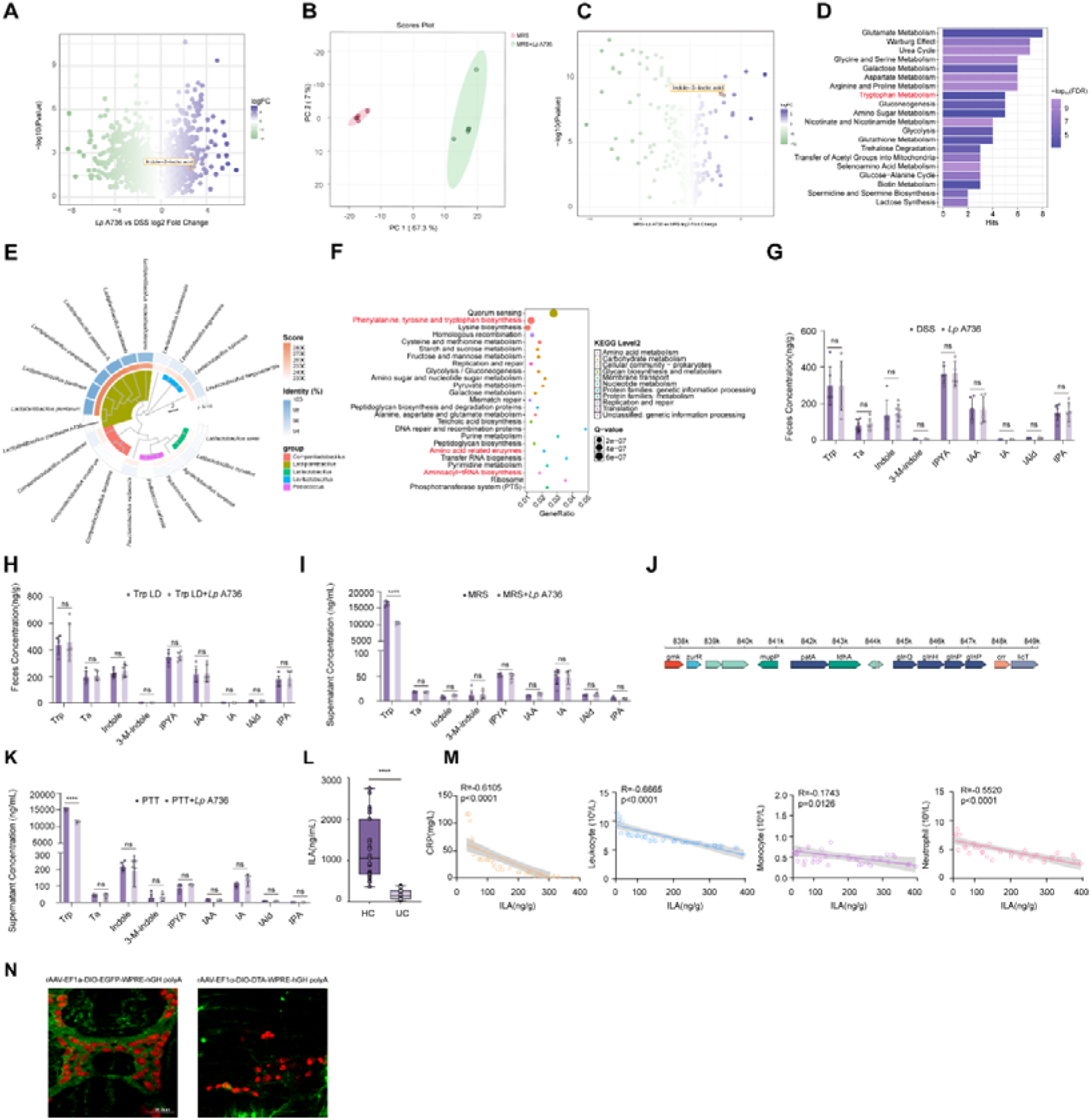
*L. plantarum* A736 impacts tryptophan metabolism and its metabolite, ILA, regulates ChAT-expressing neurons and immune responses, related to Figure 4, Figure 5 and Figure 6. (A) Volcano plot illustrating differential metabolites in feces between DSS and *Lp* A736 (n=5). (B) Principal component analysis (PCA) between MRS and MRS *+Lp* A736 by untargeted metabolome (n=5). (C) Volcano plot illustrating differential metabolites between MRS and MRS*+Lp* A736 (n=5). (D) Pathway-associated quantitative enrichment analysis (QEA) of differentially enriched metabolites between MRS and MRS *+Lp* A736 (n=5). (E) The phylogenetic trees of *Lp* A736 including average nucleotide identity (ANI). (F) KEGG pathway enrichment analysis of the *Lp* A736 genome. (G) Concentrations of tryptophan (Trp) and indole derivatives in feces from mice between DSS and *Lp* A736 (n=6). Trp, tryptophan; TA, tryptamine; Indole; 3-Methylindole; IPYA, indole-3-pyruvic acid; IAA, indole-3-acetic acid; IA, 3-indoleacrylic acid; IAld, indole-3-carboxaldehyde; IPA, Indole-3-propionic acid. (H) Concentrations of tryptophan (Trp) and indole derivatives in feces from mice treated with *L. plantarum* A736 or PBS during low-tryptophan diet between DSS and *Lp* A736 (n=6). (I) Tryptophan (Trp) and indole derivatives production by *L. plantarum* A736 in MRS medium (n=6). (J) Diagram showing the gene clusters for ILA production from *L. plantarum* A736. ldhA, a gene encoding 2-hydroxyacid dehydrogenase. (K) Tryptophan (Trp) and indole derivatives production by *L. plantarum* A736 in peptone-tryptone-tryptophan (PTT) medium (n=6). (L) ILA levels in the feces from healthy controls and UC patients (n=35). (M) Correlative analysis of ILA with clinical parameters including C-reactive protein (CRP) values, leukocyte, monocyte and neutrophil counts in UC patients (n=35). (N) ChAT (green) and HuC/D (red) immunostaining in the colonic myenteric plexus from ChAT-iCre mice injected with rAAV-EF1α-DIO-DTA-WPRE-hGH polyA or rAAV-EF1a-DIO-EGFP-WPRE-hGH polyA. Scale bar, 36.8 μm. DSS group (DSS) and *L. plantarum* A736 treatment group (*Lp* A736) and ILA treatment group (ILA) in vivo. Low-tryptophan (Trp) diet (Trp LD), low-tryptophan (Trp) diet and *L. plantarum* A736 (Trp LD+*Lp* A736). Original culture medium (Man-Rogosa-Sharpe, MRS) and *L. plantarum* A736 culture supernatant (MRS+ *Lp* A736). Peptone-tryptone-tryptophan medium (PTT) and *L. plantarum* A736 supernatants in peptone-tryptone-tryptophan medium (PTT+ *Lp* A736). PBS-treated ChAT-iCre mice infected with AAV-encoding EGFP viruses (ChAT^AAV-EGFP^), PBS-treated ChAT-iCre mice infected with AAV-encoding DTA viruses (ChAT^AAV-DTA^). Mean ± SEM shown. Data are determined by unpaired Student’s t-test (G-I and L). ns, not significant, *p < 0.05, **p < 0.01, ***p < 0.001, and ****p < 0.0001, compared with the Control group, ^#^p<0.05, ^##^p<0.01, ^###^p < 0.001, and ^####^p < 0.0001, compared with the DSS group.

